# AWmeta empowers adaptively-weighted transcriptomic meta-analysis

**DOI:** 10.1101/2025.05.06.650408

**Authors:** Yanshi Hu, Zixuan Wang, Yueming Hu, Cong Feng, Qiuyu Fang, Ming Chen

## Abstract

Transcriptomic meta-analysis enhances biological veracity and reproducibility by integrating diverse studies, yet prevailing *P*-value or effect-size integration approaches exhibit limited power to resolve subtle signatures. We present AWmeta, an adaptively-weighted framework that unifies both paradigms. Benchmarking across 35 Parkinson’s and Crohn’s disease datasets spanning diverse tissues and adaptively down-weighting underpowered studies, AWmeta yields higher-fidelity differentially expressed genes (DEGs) with markedly reduced false positives and establishes superior gene differential quantification convergence at both gene and study levels over state-of-the-art random-effects model (REM) and original studies. AWmeta requires fewer samples and DEGs from original studies to achieve substantial gene differential estimates, lowering experimental costs. We demonstrate AWmeta’s remarkable stability and robustness against external and internal perturbations. Crucially, AWmeta prioritizes disease tissue-specific mechanisms with higher functional coherence than those from REM and original studies. By bridging statistical rigor with mechanistic interpretability, AWmeta harmonizes heterogeneous transcriptomic data into actionable insights, serving as a transformative tool for precision transcriptomic integration.

## Main

The exponential expansion of publicly available transcriptomic data, propelled by high-throughput sequencing advancements^1^, presents unprecedented opportunities and concomitant challenges for uncovering robust biological insights through meta-analysis. By integrating findings across independent studies, this powerful approach transcends the limitations of individual datasets, mitigating issues of statistical power, experimental variability, tissue heterogeneity, and platform-specific biases that often obscure subtle yet pathologically relevant expression signatures^2,3^. As complex diseases increasingly defy dissection by single-study designs, meta-analysis has become indispensable for identifying reproducible biomarkers and elucidating disease pathways with enhanced confidence and precision^4,5^.

Contemporary meta-analysis methodologies predominantly fall into three primary categories^6^: *P*-value combination, effect-size integration, and rank aggregation. *P*-value-based methods, such as Fisher’s^7^, Stouffer’s Z-score^8^, and the adaptively weighted Fisher’s (AW-Fisher) technique^9^, efficiently aggregate statistical significance but typically disregard effect magnitude and directionality. Conversely, effect-size approaches quantify expression differences, with the random-effects model (REM)^10^ widely adopted due to its capacity to accommodate inter-study heterogeneity and its perceived biological interpretability^11^. REM has underpinned discoveries in diverse areas, including characterizing gut microbiome dysbiosis in Parkinson’s disease^12^, identifying predictive biomarkers for cancer immunotherapy^13^, and assessing pharmacogene expression in nonalcoholic fatty liver disease^14^. Nevertheless, REM and related models can exhibit sensitivity to outlier studies and rely heavily on assumptions about the underlying heterogeneity structure^11,15^. Rank-based techniques (e.g., RankProd and RankSum^16,17^) offer robustness against outliers but often at the cost of statistical resolution and power. Critically, these distinct methodological frameworks typically operate in isolation, failing to synergistically leverage their complementary strengths.

This methodological schism represents a fundamental constraint in transcriptomic meta-analysis, particularly impeding progress in complex disease research where robust biological inference demands both high statistical confidence in identifying dysregulated genes and accurate quantitative estimates of their expression changes^18,19^. Current *P*-value integration schemes, while adept at pinpointing consistently altered genes, offer minimal information on the magnitude or biological relevance of these alterations. Conversely, effect-size methods, though designed to quantify these changes, struggle with the pervasive heterogeneity inherent in pooling diverse experimental designs, tissue sources, or patient cohorts, potentially diminishing the estimate reliability^20^. Furthermore, existing approaches lack sophisticated mechanisms to dynamically weight studies based on intrinsic data quality or specific biological context, thereby limiting overall sensitivity and the depth of achievable insights.

To address these critical gaps, we introduce AWmeta, a novel meta-analytic framework that unifies the statistical rigor of *P*-value–based method with the quantitative power of effect-size paradigm. The core innovation of AWmeta lies in an adaptive weighting scheme that (1) identifies and up-weights the most informative studies from a heterogeneous pool, and (2) explicitly models between-dataset variability to mitigate bias arising from sample size imbalances, data sparsity, and technical heterogeneity. Validated on 35 diverse transcriptomic datasets spanning Parkinson’s and Crohn’s disease across multiple tissues, AWmeta consistently outperforms the state-of-the-art REM. It secures higher-fidelity differentially expressed gene (DEG) identification with markedly reduced false positives, exhibits greater resilience to limited sample size and DEG sparsity, and maintains remarkable stability against various perturbations. More importantly, AWmeta delivers gene differential estimates with enhanced reproducibility and biological mechanism interpretation. By reducing sample-size requirements and demonstrating robustness to noise, AWmeta empowers researchers to mine the ever-expanding corpus of public transcriptomic data more effectively, accelerating the discovery of actionable molecular insights across biomedical domains.

## Results

### Overview of AWmeta

AWmeta is a transcriptomic meta-analytical framework, uniquely synergizing statistical strengths from *P*-value and effect size integration methods (Fig. 1a and “Overview of the AWmeta framework” section in Methods). Following preprocessing of multiple transcriptomic datasets from original studies of the same disease tissue (“Transcriptomic data preprocessing” section in Methods), AWmeta implements two complementary gene-wise modules: AW-Fisher and AW-REM. The AW-Fisher module calculates meta *P*-values by optimizing study-specific weights to minimize the combined probability, effectively filtering out less informative studies while preserving statistical power. Subsequently, in AW-REM module these optimized weights are embedded into REM architecture to derive weighted fold change estimates.

**Fig. 1.**
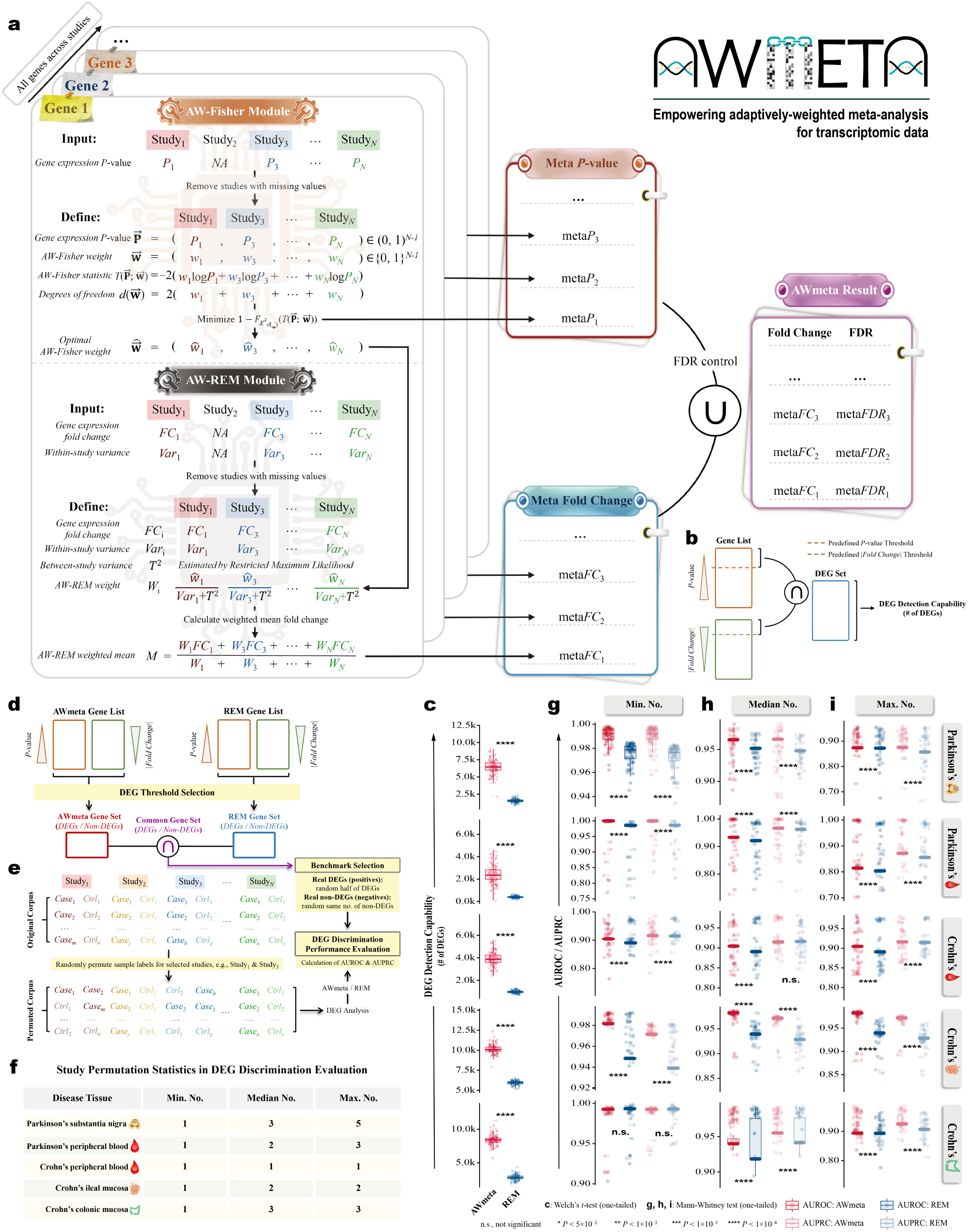
Overview of AWmeta and DEG identification evaluation. **a,** Schematic of the AWmeta framework, comprising the AW-Fisher module for *P*-value calculation and the AW-REM module for effect size (log_2_-transformed fold change) estimation. DEG identification evaluation includes DEG detection capability (**b, c**) and discrimination (**d**–**i**). **b,** Schematic defining DEG detection capability as the number of DEGs identified by predefined statistical significance (*P*-value) and gene expression fold change (log_2_-based) thresholds. **c,** DEG detection capability performance comparisons between AWmeta and REM with corrected *P*-value (FDR) < 0.01 and fold change (|log_2_FC|) > log_2_1.2 across five disease tissues. Statistical significance was determined with one-tailed Welch’s *t*-test. **d,** Strategy for generating the semi-synthetic benchmark dataset, sampling equivalent DEGs and non-DEGs from common genes identified by both AWmeta and REM. **e,** Workflow for evaluating DEG discrimination performance using sample label permutation within the semi-synthetic benchmark dataset, followed by AWmeta/REM procedure and AUROC/AUPRC calculation. **f,** Study permutation statistics (number of permuted studies) in the DEG discrimination evaluation procedure across five disease tissues. **g**–**i**, DEG discrimination performance comparisons between AWmeta and REM using minimum-(**g**), median-(**h**), and maximum-(**i**) permuted semi-synthetic simulation strategy with FDR < 0.01 and |log_2_FC| > log_2_1.2 across five disease tissues. Statistical significance was determined using one-tailed Mann-Whitney test (**g**–**i**). Textual details of the AWmeta framework, DEG detection capability, and minimum-, median-and maximum-permuted semi-synthetic simulation strategies for DEG discrimination, reside in “DEG detection capability evaluation” and “DEG discrimination evaluation using semi-synthetic simulation strategy” sections in Methods. Boxplot bounds indicate interquartile ranges (IQR), centers denote median values, and whiskers extend to 1.5 IQR. The following icons represent different tissue sources: 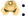 substantia nigra; 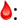 peripheral blood; 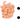 ileal mucosa; 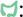 colonic mucosa.

In the following sections, we detail a multifaceted rigorous comparison of AWmeta against the state-of-the-art REM, a representative effect-size integration method^21,22^. Benchmarking on 35 transcriptomic datasets from Parkinson’s and Crohn’s disease tissues (Extended Data Fig. 1 and “Transcriptomic datasets” section in Methods), this evaluation assessed key metrics including DEG detection capability and discrimination, gene- and study-wise gene differential quantification convergence, stability and robustness, and biological relevance (“Transcriptomic meta-analysis evaluation metrics” section in Methods). AWmeta consistently demonstrated superior performance, establishing its utility as a powerful tool for precision transcriptomic integration and providing a solid foundation for downstream biological investigation.

### AWmeta secures robust higher-fidelity DEG identification across transcriptomic contexts

A primary goal of transcriptomic meta-analysis is to enhance statistical power for identifying DEGs reliably, i.e., to detect more subtle yet vital DEGs, typically defined by statistical significance and fold-change thresholds (Fig. 1b and “DEG detection capability evaluation” section in Methods). Systematically benchmarking across five distinct disease tissue contexts using nine combinations of statistical significance (0.01, 0.05, and 0.10) and fold-change thresholds (log_2_ 1.2, log_2_ 1.5, and log_2_ 2.0), AWmeta consistently identified significantly more DEGs than REM (*P* < 10^−4^, one-tailed Welch’s *t*-test over 100 bootstrap iterations; Fig. 1c and Extended Data Fig. 2). For instance, under a specific threshold combination (*P* < 0.01 and log_2_ FC *>* log_2_ 1.2), AWmeta yielded 69–475% increases in detected DEGs versus REM across all tissues (Fig. 1c), which demonstrates AWmeta’s superior statistical sensitivity in DEG detection.

A key challenge in meta-analysis is to increase statistical power while rigorously controlling false positives. To formally evaluate this trade-off, we designed a semi-synthetic simulation framework to assess DEG discrimination (“DEG discrimination evaluation using semi-synthetic simulation strategy” section in Methods). Inspired by Li and colleagues^23^, this framework first creates benchmark datasets with a known ground truth of positives (DEGs) and negatives (non-DEGs) (Fig. 1d). We then systematically challenged AWmeta’s performance by degrading the biological signal in a controlled manner. This was achieved by permuting sample labels in a progressively increasing number of studies within each tissue context (Fig. 1e). Specifically, we simulated three distinct noise scenarios by permuting the minimum, median, and maximum allowable number of studies, where DEG discrimination was quantified by the area under the receiver operating characteristic curve (AUROC) and the area under the precision-recall curve (AUPRC) over the above defined benchmark genes (Fig. 1f). This perturbation design enabled a rigorous assessment of AWmeta’s resilience across diverse data quality landscapes, a critical feature for real-world applications. Across all simulated noise levels, AWmeta consistently outperformed or was comparable to REM in DEG discrimination.

Under minimum-permuted low-noise condition, AWmeta demonstrated clear and significant advantage across nearly all tissue contexts and DEG thresholds (*P* < 10^−4^, 10^−3^, 10^−2^ or 0.05, one-tailed Mann-Whitney test; Fig. 1g and Extended Data Fig. 3). As expected, performance decayed for both methods with increasing noise from median and maximum study permutations. However, AWmeta’s superiority over REM was not only maintained but often became more pronounced under these more challenging conditions (*P* < 10^−4^, one-tailed Mann-Whitney test; Fig. 1h,i and Extended Data Fig. 4 and 5). Notably, AWmeta’s performance remained remarkably robust even under high-noise scenarios, with median AUROC and AUPRC exceeding 0.85 in most cases (Extended Data Fig. 5), highlighting its ability to effectively discount noise from potentially confounding studies. Taken together, these results demonstrate that AWmeta achieves a superior balance between heightened detection sensitivity and robust discrimination for higher-fidelity DEG identification from heterogeneous transcriptomic datasets.

### AWmeta establishes superior gene- and study-wise convergence in gene differential quantification

To rigorously assess AWmeta’s ability to synthesize a consensus biological signal from heterogeneous transcriptomic datasets, we evaluated its convergence at both gene and study levels. We first quantified gene-wise convergence—the proximity of a gene’s meta effect-size estimates to ones from original studies—using the mean absolute deviation (MAD) between meta and original fold changes (Fig. 2a and “Gene-wise convergence assessment for gene differential quantification” section in Methods), where lower MADs signify more accurate biological representations.

**Fig. 2.**
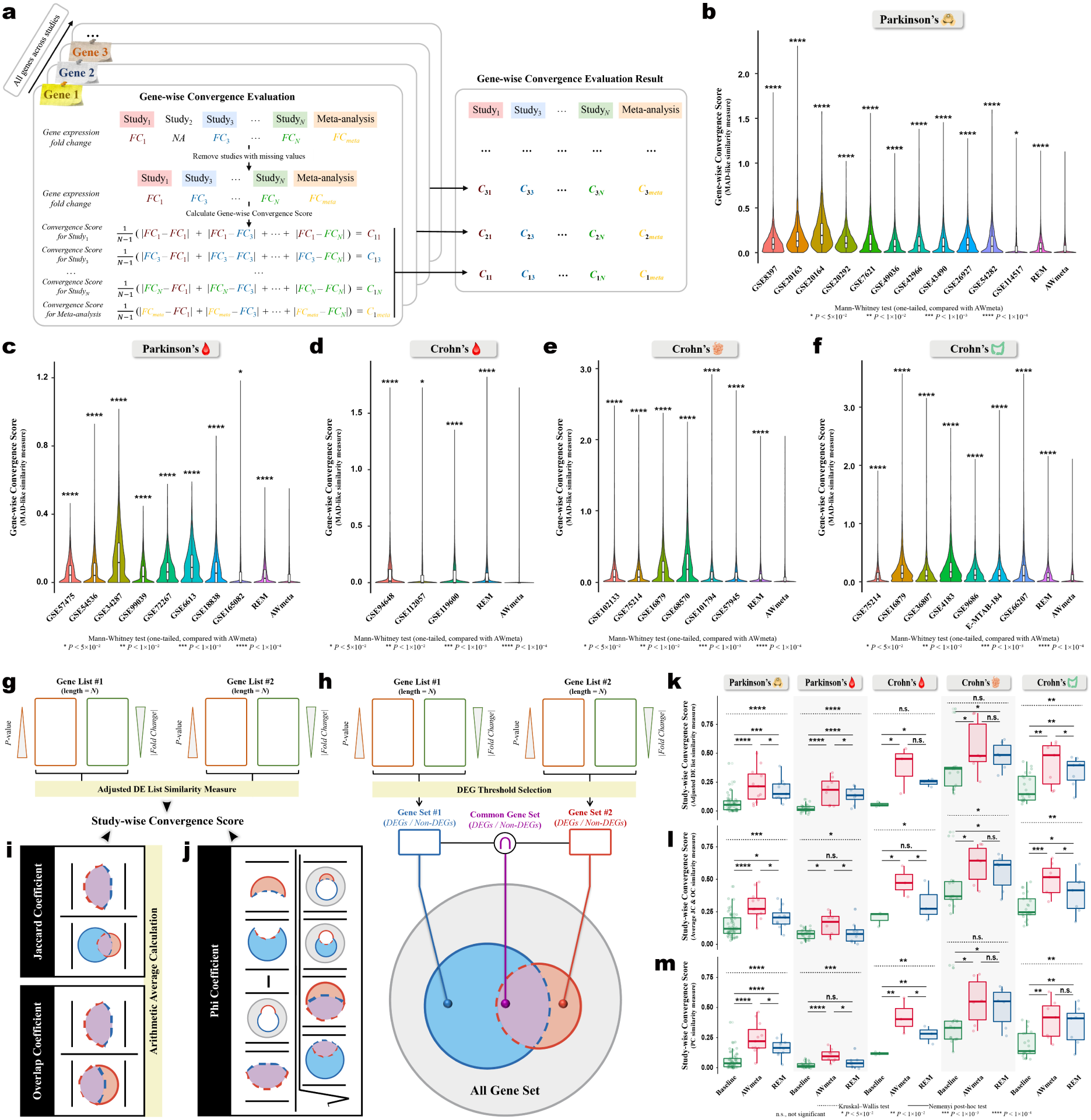
AWmeta establishes superior gene- and study-wise convergence in gene differential quantification. To benchmark meta-analysis efficacy across biological scales, we assessed gene differential quantification convergence in gene-(**a**–**f**) and study-(**g**–**m**) wise manners. **a**–**f**, For gene-wise convergence, mean absolute deviation (MAD)-like similarity measure was utilized to quantify the per-gene fold change (|log_2_FC|) similarity among AWmeta, REM and original studies (**a**), with smaller value indicating better convergence, which demonstrates AWmeta’s superior gene-wise convergence over REM and orignal studies across five disease tissues: Parkinson’s substantia nigra (**b**) and peripheral blood (**c**), and Crohn’s peripheral blood (**d**), ileal mucosa (**e**) and colonic mucosa (**f**). Statistical significance against AWmeta for gene-wise convergence comparisons was determined by one-tailed Mann-Whitney test. **g**–**m**, Three complementary similarities, i.e., adjusted DE list similarity (**g**, conceptual schematic; **k**, assessment result), the arithmetic average of Jaccard (JC) and overlap coefficient (OC) (**h, i**, conceptual schematic; **l**, assessment result) and phi coefficient (PC) (**h, j**, conceptual schematic; **m**, assessment result), were used to derive study-wise convergence score, indicating AWmeta exerts better convergence than REM and baselines in five disease tissues with FDR < 0.05 and |log_2_FC| > log_2_1.2. For comparison purpose, results from original studies serve as reference baselines. Overall study-wise convergence differences among AWmeta, REM and baselines were tested with Kruskal–Wallis test, followed by Nemenyi post-hoc test for pairwise comparisons. Detailed description for MAD-like gene-and three study-wise convergence similarity measures can be referred to in “Gene-wise convergence assessment for gene differential quantification” and “Study-wise convergence assessment for gene differential quantification” sections in Methods and Extended Data Fig. 7. Boxplot bounds show interquartile ranges (IQR), centers indicate median values, and whiskers extend to 1.5 IQR. The following icons represent different tissue sources: 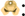 substantia nigra; 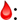 peripheral blood; 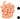 ileal mucosa; 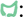 colonic mucosa.

AWmeta consistently yielded significantly lower gene-wise convergence scores in all five disease tissues, compared to both REM and baseline ones (Fig. 2b–f; *P <* 10^−4^ or 0.05, one-tailed Mann-Whitney test against AWmeta). Notably, while some original studies occasionally outperformed REM in specific contexts, AWmeta (merely 57–74% of REM) consistently achieved lower scores than any original study across all tissues, which suggests its capacity to robustly identify and integrate reliable signals while effectively down-weighting divergent studies, thereby providing a superior consensus representation of the gene expression landscape. Since disease processes are primarily driven by DEGs, we further confirmed this superior performance was still evident for these specific genes by the same assessment paradigm using nine distinct thresholds, combining three significance levels (0.01, 0.05, and 0.10) and three fold-change cutoffs (log_2_ 1.2, log_2_ 1.5, and log_2_ 2.0). Across all threshold and disease tissue scenarios, AWmeta maintained evidently lower convergence scores, accounting for 56–80% of REM (Extended Data Fig. 6a–e; *P <* 10^−4^, 10^−3^, 10^−2^ or 0.05, one-tailed Mann-Whitney test over AWmeta), which underscores AWmeta’s effectiveness to derive robust fold-change estimates for disease-relevant genes, independent of specific statistical criteria.

Next, we evaluated study-wise convergence to determine how well the meta-analytic results reflect the collective evidence across all contributing studies. We employed three complementary approaches: an adjusted rank-sensitive similarity metric emphasizing top-ranked genes (denoted “adjusted DE list similarity” thereafter), the arithmetic mean of Jaccard and overlap coefficients (JC/OC) for DEG concordance, and the phi coefficient (PC)^24^ to assess classification agreement beyond chance (Fig. 2g–j, Extended Data Fig. 7 and “Study-wise convergence assessment for gene differential quantification” section in Methods). Higher scores indicate better study-wise convergence for all metrics.

Across the five disease tissues, AWmeta consistently achieved significantly higher study-wise convergence scores than baselines representing original inter-study agreement, with dramatic 30–1,166% improvements (Fig. 2k–m; *P* < 0.05 and log_2_ FC *>* log_2_ 1.2 where applicable). While overall convergence scores tended to be lower in Parkinson’s over Crohn’s disease tissues, potentially reflecting higher inherent variability within these specific disease contexts, AWmeta significantly outperformed REM in the majority (10 out of 15) of comparisons across different metrics and tissues, performing comparably otherwise, particularly pronounced in tissues like Parkinson’s and Crohn’s peripheral blood, where AWmeta’s convergence scores improved by 35–156% compared to REM (Fig. 2k–m). To further validate these findings for the JC/OC and PC metrics, we confirmed AWmeta’s superior performance across nine different DEG cutoffs (Extended Data Fig. 8). These results indicate that the gene differential quantification results processed by AWmeta are more representative of the faithful consensus signal across studies than those derived from REM or original studies.

### AWmeta attains accelerated meta-analysis convergence with reduced samples and DEGs

While large sample sizes are known to enhance transcriptomic differential quantification efficacy, the feasibility of achieving reliable meta-analytic effect sizes from studies with limited samples has remained an open question. To fill this gap, we systematically correlated study-wise convergence with sample size and employed Spearman correlation with a two-tailed significance test to capture potentially non-linear dependencies. To ensure robustness and minimize sensitivity to arbitrary cutoffs, this association analysis leveraged results derived from nine distinct DEG thresholds (as previously described) across both JC/OC and PC metrics.

We observed pronounced positive correlations between sample size and study-wise convergence in Parkinson’s substantia nigra and peripheral blood, as well as in Crohn’s ileal mucosa, for both AWmeta and REM (Fig. 3a–c). This reinforces the principle that larger cohorts generally yield results that more closely approximate the consensus biological signal. Crucially, at equivalent sample sizes, AWmeta consistently outperformed REM in convergence across all five disease tissues, where performance gaps widened progressively with increasing sample size. To illustrate, using the adjusted DE list similarity metric in Crohn’s ileal mucosa, while AWmeta and REM exhibited comparable convergence scores (~ 0.38) at the same sample size of 62, AWmeta’s score surged to 0.77 at a sample size of 200—a 54% increase over REM’s 0.50 (Fig. 3a), which demonstrates AWmeta can reach much better convergence level under the same sample-size context. Notably, AWmeta attains comparable convergence with considerably fewer samples relative to REM. To reach a convergence score of 0.2 in Parkinson’s substantia nigra, for instance, AWmeta necessitated 44% or 43% fewer samples than REM with JC/OC (15 versus 27; Fig. 3b) or PC metric (20 versus 35; Fig. 3c). This marked improvement in sample efficiency translates directly into substantial reductions in experimental cost and resource allocation.

**Fig. 3.**
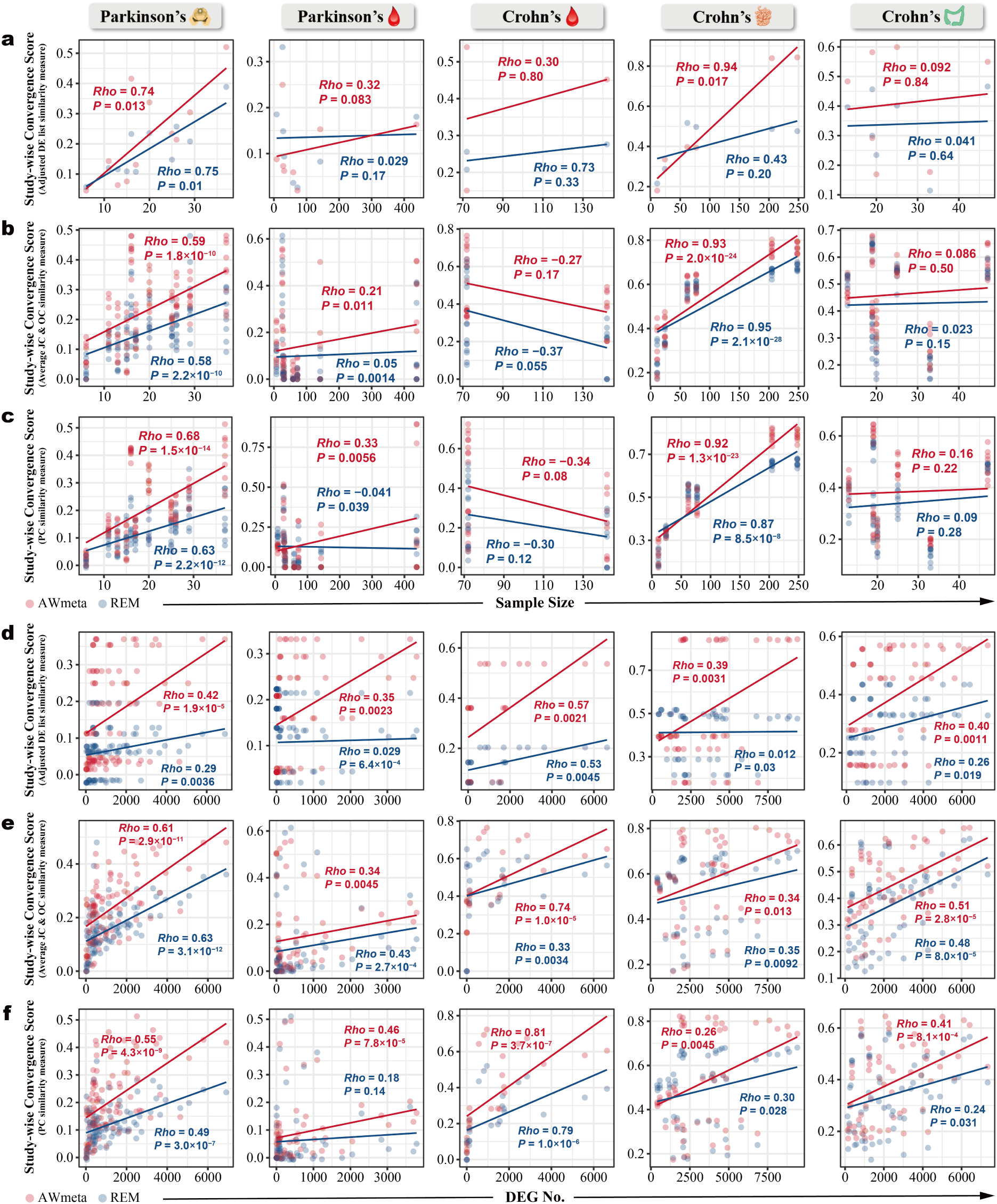
AWmeta attains accelerated meta-analysis convergence with reduced samples and DEGs. **a**–**f**, We dissected the correlations of per-study sample size and DEG number with study-wise convergence measured by three complementary similarities, i.e., adjusted DE list similarity (**a, d**), the arithmetic average of Jaccard (JC) and overlap coefficient (OC) (**b, e**) and phi coefficient (PC) (**c, f**) across five disease tissues, showcasing AWmeta’s accelerated study-wise meta-analysis convergence against REM with fewer samples and DEGs, in which non-linear correlations were quantified by spearman’s rho with two-tailed significance test. To enhance statistical power and mitigate threshold sensitivity, this correlation analysis summarized results from nine different DEG threshold combinations, spanning varying significance levels (0.01, 0.05 and 0.10) and fold change cutoffs (log_2_1.2, log_2_1.5 and log_2_2.0), for both the average of JC and OC and PC similarity measures. Detailed description for these three study-wise convergence similarity measures can be referred to in “Study-wise convergence assessment for gene differential quantification” section in Methods, Fig. 2g–j and Extended Data Fig. 7. The following icons represent different tissue sources: 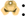 substantia nigra; 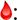 peripheral blood; 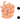 ileal mucosa; 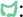 colonic mucosa.

We further hypothesized that DEG abundance would positively contribute to accurate meta-analytic estimates for gene differential quantification and substantiated this premise through the observation of robust positive correlations between DEG number and study-wise convergence across all five disease tissues (Fig. 3d–f), following the analogous procedure to the above sample-size correlation analysis. For studies with similar DEG counts, AWmeta constantly yielded higher convergence scores than REM, with performance disparity more pronounced as DEG abundance ascended, suggesting AWmeta possesses a heightened sensitivity to the underlying biological signal embedded within scattered DEGs. For example, to arrive at a PC metric-based convergence score of 0.4 in Crohn’s peripheral blood, AWmeta required only 2,000 DEGs, whereas REM exacted approximately 4,600—a 130% escalation—for equivalent resolution (Fig. 3f). This highlights AWmeta’s capacity to achieve robust convergence even from datasets with sparser DEG profiles, a practical advantage for experimental designs where DEG discovery may be limited.

Collectively, these findings imply AWmeta can deliver reliable meta-analytic outcomes in less stringent experimental scenarios, such as studies involving milder treatments or designs with fewer replicates, which substantially mitigates experimental complexity and cost, especially for ambitious large-scale research programs.

### AWmeta delivers remarkable stability and robustness in transcriptomic integration

We sought to determine whether AWmeta’s adaptively-weighted strategy confers superior stability and robustness to gene differential meta-estimates against REM, and designed quantitative metrics to evaluate consistency over random splits and resilience to systematic perturbations across the five disease tissues.

First, we assessed stability by quantifying the concordance of ranked gene differential lists derived from randomly halved sample sets within each study, a process replicated across 100 iterations (Fig. 4a and “Stability and robustness assessment of transcriptomic integration” in Methods). Across all five disease tissues, AWmeta exhibited markedly higher stability scores relative to REM (Fig. 4b; *P* < 10^−4^, one-tailed Welch’s *t*-test), underscoring its enhanced consistency under data rationing. The observation that median stability scores for both methods were below 0.7, is likely attributable to the inherently reduced statistical power and study-wise convergence that accompanies halving the sample size (Fig. 3a).

**Fig. 4.**
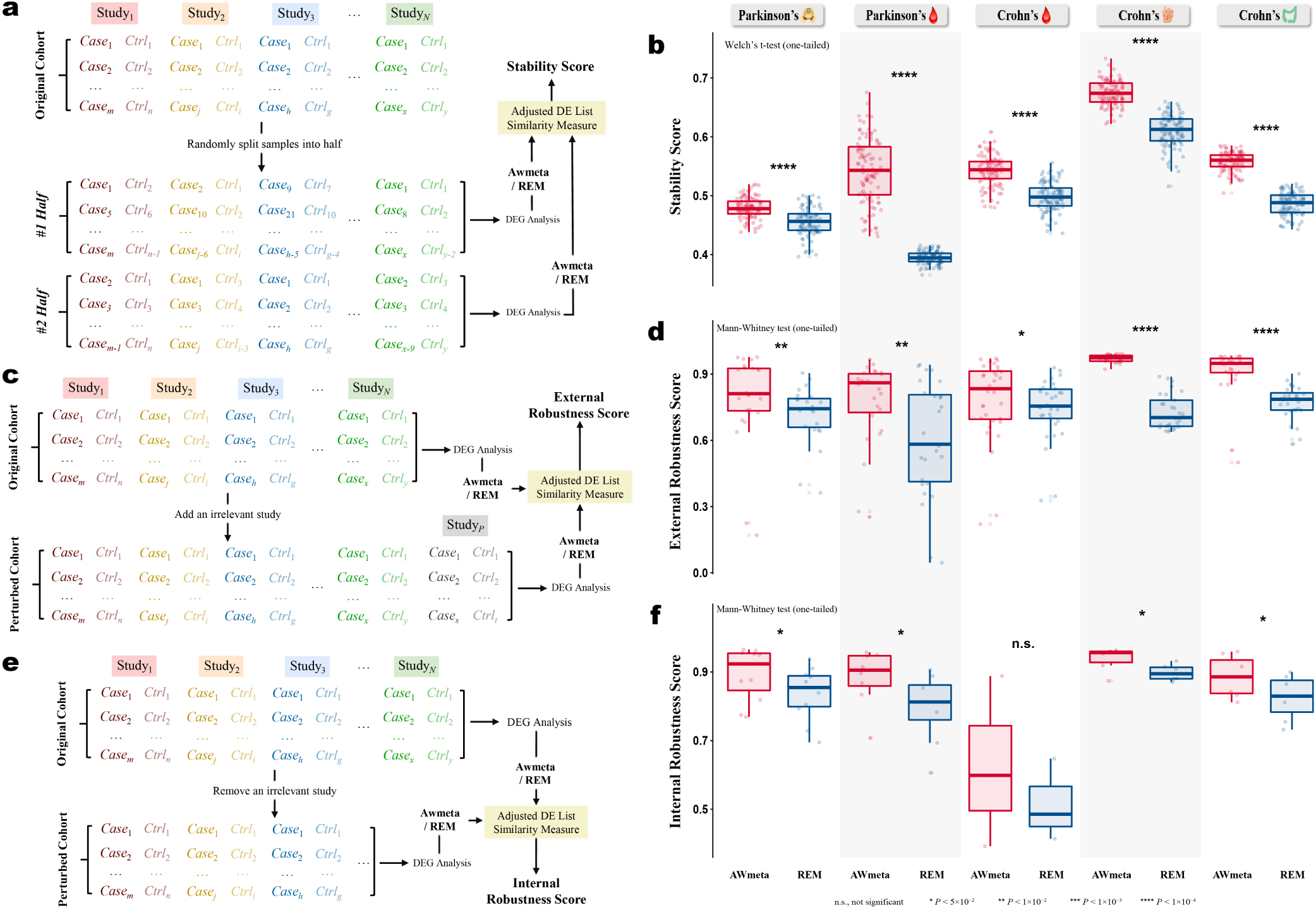
AWmeta delivers remarkable stability and robustness in transcriptomic integration. **a,** Workflow for evaluating the stability of transcriptomic integration. **b,** Stability assessment against AWmeta and REM with one-tailed Welch’s *t*-test. **c**–**f**, We assessed robustness against external interference (**c,** conceptual schematic; **d,** assessment result) and internal defects (**e,** conceptual schematic; **f,** assessment result) in transcriptomic integration across five disease tissues, with one-tailed Mann-Whitney test, indicating AWmeta’s superior performance over REM. Detailed description of adjusted DE list similarity measure can be referred to in “Study-wise convergence assessment for gene differential quantification” section in Methods and Extended Data Fig. 7. Boxplot bounds show interquartile ranges (IQR), centers indicate median values, and whiskers extend to 1.5 IQR. The following icons represent different tissue sources: 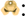 substantia nigra; 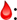 peripheral blood; 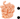 ileal mucosa; 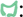 colonic mucosa.

We then challenged the robustness of each method against two distinct forms of perturbation: external interference, simulated by the inclusion of a thematically unrelated study (Fig. 4c and “Stability and robustness assessment of transcriptomic integration” in Methods), and internal fragility, evaluated through a systematic leave-one-study-out procedure (Fig. 4e and “Stability and robustness assessment of transcriptomic integration” in Methods). Against external interference, AWmeta displayed remarkable resilience with median robustness scores above 0.8 and established a significant performance margin over REM across all tissues (Fig. 4d; *P* < 10^−4^, 10^−2^ or 0.05, one-tailed Mann-Whitney test). This capacity to resist discordant data is a direct consequence of AWmeta’s adaptive weighting scheme, which effectively minimizes the influence of outlier studies.

In internal robustness assessment, AWmeta again achieved significantly higher scores than REM in four of the five tissues (Fig. 4f; *P* < 0.05, one-tailed Mann-Whitney test). The sole exception was Crohn’s peripheral blood, where the small cohort of only three studies constrained the median robustness scores to below 0.6 for both methods. Notably, in tissues comprising six or more studies, AWmeta achieved exceptional median internal robustness scores around 0.9, demonstrating highly consistent results even upon the exclusion of individual constituent studies.

These rigorous stress tests validate that AWmeta’s adaptive weighting architecture endows the meta-analytic process with significantly strengthened stability and robustness. This reinforcement ensures the derivation of more dependable biological insights when integrating diverse and inherently heterogeneous transcriptomic datasets.

### AWmeta facilitates discovery of disease-relevant genes

A pivotal determinant of a meta-analysis method’s utility is its capacity to prioritize genes of genuine pathological importance. To rigorously assess this, we quantified the biological relevance of gene rankings from AWmeta, REM, and the original studies (as baselines) against authoritative Parkinson’s and Crohn’s disease-gene benchmarks—compiled from DisGeNET^25^, MalaCards^26^, and an in-house curated genetic variation corpus^27^—using a custom metric that integrates both statistical significance and effect size magnitude, which provides an objective and threshold-agnostic evaluation of gene prioritization performance (Fig. 5a and “Biological relevance assessment of gene differential quantification” section in Methods).

**Fig. 5.**
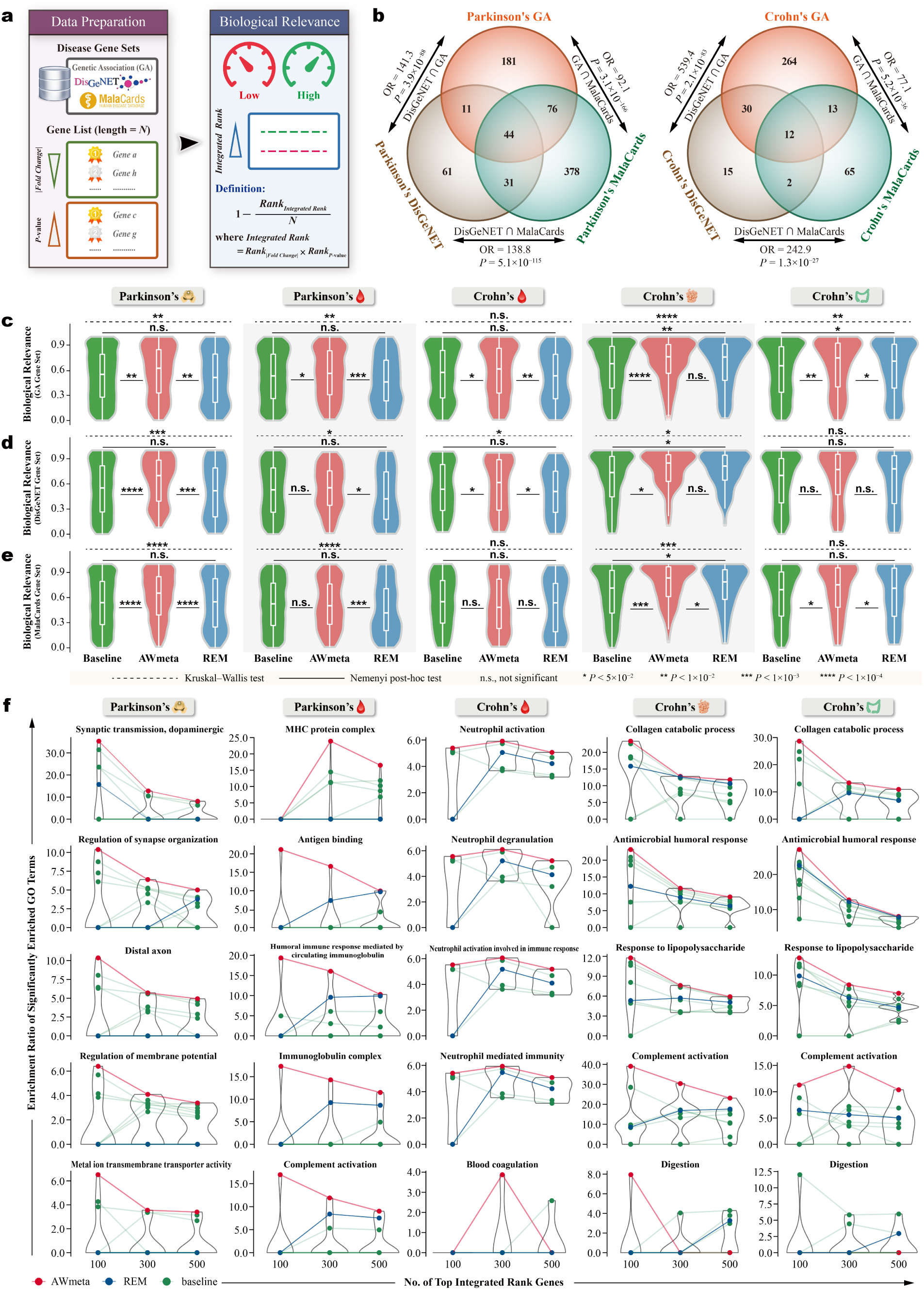
AWmeta enhances identification and mechanism interpretation of disease tissue-specific genes. **a,** Workflow for quantifying gene-wise biological relevance against three benchmark gene sets of Parkinson’s and Crohn’s disease from genetic association (GA) variation corpus, DisGeNET and MalaCards, with higher scores indicating stronger tissue-contextual disease associations. **b,** Pairwise coherence analysis of the three benchmark gene sets for Parkinson’s and Crohn’s disease. The degree of overlap between benchmarks was quantified using odds ratios (OR), with statistical significance determined by Fisher’s exact test. **c**–**e**, Biological relevance evaluations on AWmeta (red), REM (blue) and baselines (green) against GA (**c**), DisGeNET (**d**) and MalaCards (**e**) benchmarks, where higher scores (y-axis) denote enhanced biological relevance. Original study-derived biological relevance results serve as reference baselines. We assessed overall differences in biological relevance scores using Kruskal-Wallis test, with Nemenyi post-hoc test for pairwise comparisons (AWmeta versus REM, AWmeta versus baseline, and REM versus baseline). Boxplot bounds indicate interquartile ranges (IQR), centers denote median values, and whiskers extend to 1.5 IQR. **f,** Dynamic GO enrichment trajectories across top-ranked (100, 300 and 500) genes identified by AWmeta (red), REM (blue) and baselines (green), with higher enrichment ratio (y-axis) indicating stronger disease-tissue involvement. Original study-derived enrichments serve as baselines. Connected lines visualize trajectory patterns across gene rank thresholds (x-axis). Textual details of the biological relevance assessment framework and detailed definition of enrichment ratio, reside in “Biological relevance assessment of gene differential quantification” section in Methods and “AWmeta enables disease tissue-specific mechanism interpretation” section in Results. The following icons represent different tissue sources: 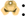 substantia nigra; 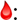 Peripheral blood; 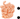 ileal mucosa; 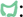 colonic mucosa.

Prior to assessing performance, we first validated the coherence of our benchmark gene sets, with overlap magnitude quantified using odds ratio (OR) and statistical significance determined by Fisher’s exact test. Pairwise comparisons revealed substantial overlaps among the three independent sources for both Parkinson’s disease (e.g., DisGeNET versus MalaCards, OR = 138.8, *P* = 5.1 × 10^−115^) and Crohn’s disease (e.g., DisGeNET versus MalaCards, OR = 242.9, *P* = 1.3 × 10^−27^) (Fig. 5b). This strong reciprocal consistency affirmed their utility for a reliable evaluation of biological relevance.

Our primary analysis revealed that AWmeta consistently generates more biologically meaningful gene rankings than REM and the baseline studies (*P* < 10^−4^, 10^−3^, 10^−2^ or 0.05, Nemenyi post-hoc test; Fig. 5c–e). Specifically, when benchmarked against our genetic variant corpus, AWmeta achieved significantly higher relevance scores across all interrogated tissues (Fig. 5c). This superior performance extended to the DisGeNET benchmark in critical disease tissues, including Parkinson’s substantia nigra and Crohn’s peripheral blood and ileal mucosa (Fig. 5d). A similar advantage was observed using the MalaCards benchmark for Parkinson’s substantia nigra and Crohn’s ileal and colonic mucosa (Fig. 5e). Notably, AWmeta’s superiority was particularly pronounced in the primary disease-affected tissues—Parkinson’s substantia nigra and Crohn’s ileal mucosa—where it surpassed baselines across all three independent benchmarks. Cumulatively, in 11 instances of the 15 tissue-benchmark comparisons (5 tissues × 3 benchmarks), AWmeta’s scores were significantly higher than those of both the baselines and REM. In stark contrast, REM failed to offer a significant improvement over baseline scores in 11 of 15 comparisons, underscoring its limited ability to distill new biological insights from existing data.

These results establish that AWmeta’s gene prioritization is not merely a statistical refinement but a substantive improvement in biological fidelity. By more effectively elevating established disease-associated genes to the top of significance rankings, AWmeta provides a clearer and more accurate representation of the underlying pathology. This enhanced resolution positions AWmeta as a powerful discovery engine, capable of transforming heterogeneous transcriptomic datasets into a focused and mechanistically coherent view of disease processes.

### AWmeta enables disease tissue-specific mechanism interpretation

We further implemented Gene Ontology (GO) enrichment to explore disease mechanism interpretation based on meta-analysis prioritized genes. To avoid arbitrariness, three thresholds (100, 300, and 500) were used to select the number of top integrated rank genes (“Biological relevance assessment of gene differential quantification” section in Methods). The enrichment ratio quantifies the degree to which GO terms are significantly enriched in relevant disease tissues:

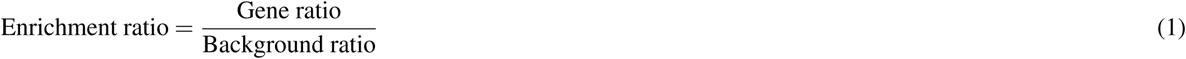

where gene ratio is the proportion of genes annotated to a specific GO term within the top integrated rank genes, and background ratio represents the analogous fraction across a reference gene set. GO terms with higher enrichment ratios are more likely to be involved in a given disease tissue. For comparison, GO enrichments derived from original studies served as baselines.

Compared with REM and baselines, representative GO terms enriched in AWmeta-derived top integrated rank genes consistently exhibited the highest enrichment ratios across nearly all five disease tissues (Fig. 5f), demonstrating AWmeta’s enhanced capacity for disease-relevant gene prioritization in tissue-specific contexts. In contrast, REM underperformed relative to some baselines (Fig. 5f), reflecting diminished biological relevance within its gene sets. For instance, biological processes related to synaptic organization and transmission (“synaptic transmission, dopaminergic”, “regulation of synapse organization”, and “distal axon”) were significantly enriched in Parkinson’s substantia nigra, consistent with their known involvement in Parkinson’s pathogenesis^28,29^. Likewise, “metal ion transmembrane transporter activity” and “regulation of membrane potential” were significantly enriched, highlighting their pivotal roles in Parkinson’s substantia nigra-involved mechanisms^30–32^. AWmeta achieved the highest enrichment for these GO terms, a trend robust across all thresholds. The enrichment ratios for AWmeta and some baselines monotonically decreased with more integrated rank genes included, indicating these term-related genes are concentrated at the very top of the ranked lists. Strikingly, REM-prioritized genes exhibited minimal or absent enrichment across all five representative GO terms, further underscoring its impaired capacity to capture contextual biological functions.

Given the well-established inflammatory pathogenesis of Parkinson’s and Crohn’s disease in non-hematopoietic tissues^33,34^, we hypothesized that blood-derived gene signatures would reflect systemic immune dysregulation and vascular barrier impairment at disease-relevant interfaces: Parkinson’s blood-brain barrier and Crohn’s intestinal vasculature. Peripheral blood analyses revealed significant enrichment of immune-related GO terms in Parkinson’s disease, including “MHC protein complex”^35^, “antigen binding”^36^, and “immunoglobulin complex”. The circulatory specificity was further evidenced by “humoral immune response mediated by circulating immunoglobulin”. Similarly, in Crohn’s peripheral blood, neutrophil-related GO terms (“neutrophil degranulation”, “neutrophil activation involved in immune response”, “neutrophil mediated immunity”, and “neutrophil activation”) indicated the involvement of immune-inflammatory processes^37^. Furthermore, significant Parkinson’s “complement activation” and Crohn’s “blood coagulation” provided disease-specific vascular insights. Aberrant complement system activity may imply blood-brain barrier disruption in Parkinson’s patients^38,39^, whereas increased venous thromboembolism risk in Crohn’s patients due to abnormal coagulation^40^ indicates intestinal vascular barrier impairment^41^.

The ileal and colonic mucosa constitute primary pathological sites in Crohn’s disease^34^, where collagen plays a key role in extracellular matrix remodeling^42^, evidenced by the significant “collagen catabolic process”. Gut microbiota dysbiosis, reflected by GO terms “antimicrobial humoral response” and “response to lipopolysaccharide”, further aligned with Crohn’s pathogenesis^43,44^. While Crohn’s ileal and colonic mucosa share multifaceted similarities, they displayed two key distinctions: complement activation and digestion. Although both mucosal tissues showed complement activation, its enrichment ratios were 2-4 times higher in ileal versus colonic mucosa, supported by immunofluorescence staining and single-cell transcriptomics indicating more active complement activation in ileal mucosa^45,46^. It has been generally acknowledged the ileal mucosa uniquely mediates hydrolase-driven enzymatic digestion, whereas the colonic mucosa plays no substantive role in chemical digestion^47^. This functional dichotomy was corroborated by the enrichments of the “digestion” GO term: AWmeta specifically detected “digestion” within top 100 ileal mucosa genes (highest enrichment), whereas REM and baselines showed delayed identification (top 300/500 genes; lower enrichment); conversely, AWmeta consistently excluded “digestion” from colonic mucosa enrichments, in contrast to sporadic false positives by these counterparts. These functional stratifications demonstrate AWmeta’s enhanced biological fidelity in resolving disease tissue-contextual gene functions.

## Discussion

Transcriptomic meta-analysis is pivotal for distilling robust biological insights from heterogeneous gene expression studies; yet, existing frameworks remain confined to either *P*-value combination or effect-size integration, imposing a trade-off between statistical sensitivity and quantitative fidelity. A unified strategy that seamlessly integrates both paradigms—capitalizing on their complementary strengths while circumventing their individual limitations—has therefore been a long-standing, unmet imperative.

AWmeta represents the first successful integration of *P*-value and effect-size aggregation methodologies in transcriptomic meta-analysis. The core innovation—a cross-module information transfer where optimized weights from *P*-value calculations directly enhance effect size estimation—effectively addresses between-study heterogeneity while maximizing consistent biological signal extraction. Indeed, the substantial variability often observed between studies, visually apparent in metrics like gene-wise convergence (Fig. 2b–f with per-study skewed distributions), highlights the prevalence of such heterogeneity and strongly supports the usage of random-effects-like frameworks such as AWmeta and REM over simpler fixed-effects models^48^. In our comprehensive evaluation across 35 datasets from Parkinson’s and Crohn’s disease, AWmeta demonstrated superior high-fidelity DEG detection that remained robust under substantial experimental noise (Fig. 1c,g–i and Extended Data Fig. 2–5). This enhanced discrimination capacity enabled identification of subtle yet biologically meaningful expression changes that conventional methods frequently miss, substantially improving the reliability and reproducibility of transcriptomic discoveries. Our convergence metrics revealed AWmeta’s practical advantages in approximating theoretical true values at both gene and study levels. It’s noteworthy that in our gene-wise convergence assessments, some original studies occasionally outperformed standard REM, even without larger sample sizes (Fig. 2b–f and Extended Data Fig. 6). While not conclusive, this hints that inherent study quality or specific experimental contexts might significantly influence reliability, perhaps as much as sample size itself. We also observed a tendency for these well-performing studies to utilize RNA-seq technology. These observations underscore the complexity of integrating diverse datasets and highlight the benefit of AWmeta’s adaptive capability which weights studies based on informational content rather than relying solely on metrics like sample size. AWmeta achieved equivalent study-wise convergence with significantly fewer samples and DEGs than conventional methods—a critical advantage in resource-constrained research environments that can substantially reduce experimental costs and researcher workload.

The superior biological relevance of AWmeta’s findings was rigorously established through two orthogonal and comple-mentary assessment paradigms: (1) Using authoritative disease-specific gene sets from DisGeNET, MalaCards, and in-house genetic variant corpus, AWmeta demonstrated significantly enhanced biological meaningfulness. In 11 of 15 tissue-benchmark combinations, AWmeta outperformed both REM and original studies in biological relevance scoring (Fig. 5c–e). This consistent advantage provides researchers with more accurate representations of core disease pathways and creates unprecedented oppor-tunities for discovering novel pathophysiological relationships that remain obscured in conventional analyses. (2) Longitudinal tracking of GO term enrichment across gene rank thresholds revealed AWmeta’s unique capacity to concentrate functionally critical genes within leading ranks. While terms of secondary importance (e.g., “MHC protein complex”, “blood coagulation”) showed delayed enrichment beyond top 300 ranks across all methods (Fig. 5f), pathologically central functions exhibited exclusive early enrichment in AWmeta. Crucially, terms like “digestion” in Crohn’s ileal mucosa reached peak enrichment exclusively within AWmeta’s top 100 genes (Fig. 5f), with no detection at expanded thresholds; REM failed to detect this pivotal function at both top 100 and 300 thresholds, achieving only marginal detection at top 500 (Fig. 5f)—demonstrating its fundamental limitations in biological resolution. This enrichment trajectory analysis establishes a dual-purpose paradigm for quantitatively evaluating gene prioritization performance and objectively stratifying biological mechanisms by pathological centrality. Together, these orthogonal validation strategies—leveraging curated knowledgebases and temporal enrichment dynamics—provide compelling evidence that AWmeta uniquely reconciles statistical rigor with biological fidelity, transforming heterogeneous transcriptomic data into precisely stratified mechanistic insights.

While AWmeta represents a significant advance, several aspects warrant consideration for broader application. The method’s computational complexity, though tractable for typical multi-study analyses, may require optimization for emerging consortia-level datasets exceeding 100 studies. Performance is also intrinsically linked to input data quality; while adaptive weighting mitigates variable study quality (Fig. 2b–f and Extended Data Fig. 6), unaddressed technical artifacts (e.g., severe batch effects) could subtly influence results. Furthermore, the current framework focuses on gene-level differential quantification; extension to isoform-resolution or splicing analysis would require adaptation to handle increased dimensionality. Finally, scenarios involving extreme, systematic confounding across studies (e.g., irreconcilable patient stratification) remain challenging—a limitation pervasive among meta-analytic methods.

These considerations highlight clear pathways for AWmeta’s evolution, complementary to its core strengths. Algorithmic refinements such as distributed computing could enhance scalability for ultra-large-scale integrations. Furthermore, incorporating tissue-specific molecular networks or multi-omic layers (e.g., epigenomics, proteomics) would refine biological inference beyond expression-centric views. Notably, coupling AWmeta with pharmacological databases holds significant promise for *in silico* drug target prioritization and biomarker discovery. The framework also uniquely empowers researchers to integrate limited-scale local datasets with public repositories, democratizing access to robust meta-analysis and amplifying statistical power for domain investigations.

Crucially, the present AWmeta implementation already delivers immediate, high-impact utility. It establishes a robust new standard for extracting reliable biological signals from complex, heterogeneous transcriptomic data—overcoming a fundamental methodological dichotomy that has long constrained the field. By synergistically combining *P*-value and effect size paradigms, AWmeta enhances reproducibility and biological interpretability, directly accelerating the translation of transcriptomic discoveries into clinical insights and biotechnological applications. This methodological leap provides an integral tool for navigating the growing complexity of precision transcriptomic integration in biomedical research.

## Methods

### Overview of the AWmeta framework

The transformative potential of AWmeta stems from its adaptively weighting scheme, which strategically prioritizes the most informative studies while robustly mitigating noise and outliers to yield biologically coherent, high-fidelity meta-analytic estimates. This framework performs gene-by-gene meta-analysis of heterogeneous transcriptomic studies by integrating per-study gene summary statistics (Fig. 1a). For each gene, input from each study in the valid set (*S_gene_*) comprises: (1) a *P*-value (*P_i_*), (2) a log_2_-based fold change (*FC_i_*), and (3) its corresponding within-study variance (*Var_i_*), all derived from the original gene differential quantification analyses. AWmeta consists of two sequential modules: *AW-Fisher* for adaptive *P*-value aggregation and *AW-REM* for adaptive effect-size integration. Studies absent from *S_gene_* (e.g., Study_2_ with missing *P*_2_, *FC*_2_, or *Var*_2_) are excluded a priori.

### AW-Fisher module (adaptive P-value integration)

Within this module, each gene’s meta *P*-value is obtained by selecting an optimal subset of *S_gene_* that minimizes a weighted Fisher’s statistic-derived combined *P*-value^9^. Let *N*^′^ = |*S_gene_*| be the number of studies reporting *P*-values for the gene, with S′*_gene_* = {1*,…, N*^′^} enumerating the study indices, and denote their *P*-values by 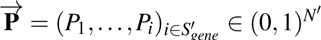. The corresponding binary weight vector, ^→−^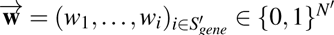, indicates inclusion (*w_i_* = 1) or exclusion (*w_i_* = 0) of Study*_i_* ∈ *S’_gene_* in the final subset. The AW-Fisher statistic is defined as:

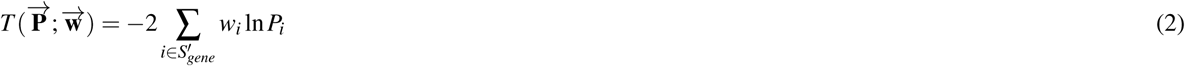

The significance level of *T* 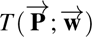 under the null hypothesis is calculated using the chi-squared distribution:

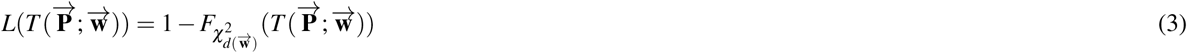

where the degrees of freedom are 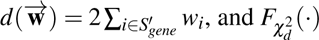 is the cumulative distribution function of the chi-squared distribution with *d* degrees of freedom.

The meta *P*-value, 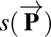, is the minimum significance level obtained by optimizing the weight vector over the studies in *S_gene_*:

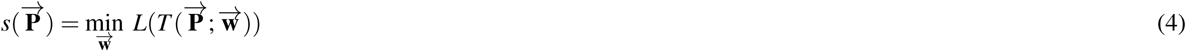

The optimal weight vector 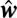 that achieves this minimum is determined by:

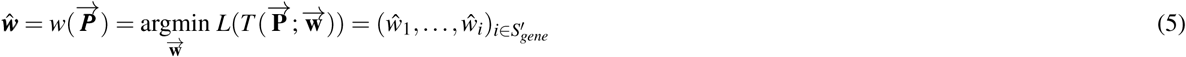

This optimal weight vector 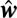, containing binary weights for each study in *S_gene_*, is passed to the following AW-REM module.

### AW-REM module (adaptive effect-size integration)

This module calculates the meta effect size (log_2_FC) using an adaptively-weighted REM. It leverages the log_2_FC (*FC_i_*) and within-study variance (*Var_i_*) from studies in *S_gene_*, modulated by the optimal binary weights 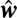 derived from the AW-Fisher module for those same studies. The contribution weight for Study*_i_* ∈ *S_gene_* in AW-REM is defined as:

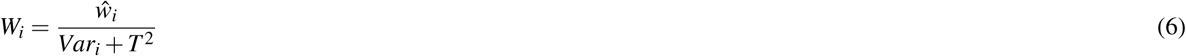

where 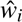 is the binary weight (0 or 1) for Study*_i_* Sgene from Eq. 5. *Var_i_* is the within-study variance for the gene in ∈ Study*_i_*, and *T* ^2^ represents the between-study variance, estimated using restricted maximum likelihood (REML) method^49^. *W_i_*is zero whenever 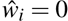, thus automatically omitting studies not selected by AW-Fisher module.

The final meta fold change, denoted **M**, is computed as an adaptively calibrated average of the study-wise effect sizes:

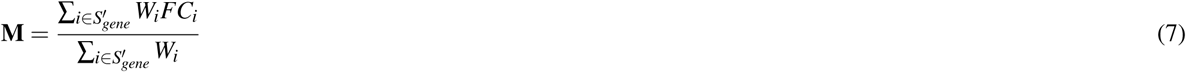

This formulation delivers a consensus fold-change estimate both statistically rigorous and quantitatively faithful to the most informative subsets of heterogeneous studies.

### Transcriptomic datasets

To proof-of-concept the AWmeta framework, we compiled 35 publicly available human transcriptomic datasets for Parkinson’s and Crohn’s disease from the Gene Expression Omnibus (GEO)^50^, Sequence Read Archive (SRA)^51^, and ArrayExpress^52^. These datasets, encompassing both microarray and RNA-sequencing (RNA-seq) platforms, included samples derived from Parkinson’s substantia nigra^53–60^ and peripheral blood^56,61–68^, Crohn’s peripheral blood^69–71^, ileal mucosa^72–77^ and colonic mucosa^76–82^. A complete list of the datasets, detailing data accession IDs, sequencing platform identifiers, dataset and tissue sources, and patient and control sample sizes, is provided in Extended Data Fig. 1.

### Transcriptomic data preprocessing

Due to the inclusion of datasets generated on different platforms, specific preprocessing pipelines were applied separately to microarray and RNA-seq data.

### Microarray data processing

To ensure accurate and up-to-date probe annotations, microarray probe identifiers were mapped to Entrez Gene IDs using information retrieved from GEO SOFT files, platform-specific Bioconductor annotation packages, and the AnnoProbe R package. For genes represented by multiple probes, we retained the probe with the largest interquartile range (IQR) of intensities across samples to maximize biological informativeness^21,83^. Subsequently, the limma R package^84^ was utilized for microarray data preprocessing, normalization, and differential gene identification. Gene differential quantification (case versus control) within each study was determined via empirical Bayes moderated *t*-statistics, yielding per-gene *P_i_*, *FC_i_*, and *Var_i_*.

### RNA-seq data processing

RNA-seq data were processed through an automated snakemake workflow^85^. Raw sequencing reads were processed with Trimmomatic^86^ to remove adapter sequences and low-quality bases. Following best practice recommendations^87^, the cleaned reads were aligned to the human reference genome (GRCh38 assembly) using HISAT2^88^. Gene-level read counts were quantified from the aligned reads using featureCounts^89^. Finally, we performed gene differential quantification with DESeq2^90^, producing per-gene *P_i_*, *FC_i_*, and *Var_i_* for each study.

### Transcriptomic meta-analysis evaluation metrics

To impartially evaluate AWmeta’s performance advances, we conducted a multi-dimensional comparison against the current gold-standard REM method^6,21,22^ across the following critical analytical domains: (i) DEG detection capability, (ii) DEG discrimination, (iii) gene- and study-wise gene differential quantification convergence, (iv) stability and robustness, and (v) biological relevance. Both methods operated on matching inputs and identical gene sets, ensuring an equitable performance assessment.

### DEG detection capability evaluation

DEG detection capability is defined as the gene count satisfying pre-defined thresholds for both corrected statistical significance *P*-value (FDR) and fold change magnitude (|log_2_FC|) (Fig. 1b). To assess the stability and reliability of this capability, we implemented a bootstrap resampling strategy with 100 iterations. In each iteration, we created bootstrapped datasets by randomly sampling with replacement from the original case and control groups while maintaining the original sample sizes, followed by meta-analysis. The resulting DEG counts formed a distribution for statistical comparison with one-tailed Welch’s *t*-test.

### DEG discrimination evaluation using semi-synthetic simulation strategy

To evaluate the ability to discriminate between DEGs and non-DEGs, particularly considering potential false positives arising from higher detection sensitivity, we adopted an evaluation metric based on semi-synthetic simulated data, inspired by Li and colleagues^23^. This approach consisted of benchmark dataset generation and evaluation using datasets with simulated noise (Fig. 1d–f) and for each tissue context:

1. Identify the intersection of DEGs and non-DEGs called by both AWmeta and REM under predefined screening thresholds (Fig. 1d).
2. Randomly sample half of the intersected DEGs to form an unbiased positive benchmark; sample an equal-sized negative benchmark from the intersected non-DEGs (Fig. 1d).
3. Construct semi-synthetic datasets by permuting case/control labels within a subset of original studies (e.g., Study_1_ and Study_3_; Fig. 1e). Label permutation removes true signal from those studies.
4. Apply AWmeta and REM to the combined set of original and label-permuted studies; compute the AUROC and AUPRC over the previously defined positive and negative benchmark genes (Fig. 1e).
5. Repeat Steps 3 and 4 for 100 times to obtain distributions of AUROC and AUPRC, summarizing performance under minimum-, median-, and maximum-permuted scenarios (Fig. 1f), which ensures the stability and reliability of our assessment. Statistical significance between AWmeta and REM was tested via one-tailed Mann–Whitney test.

### Gene-wise convergence assessment for gene differential quantification

To assess per-gene differential quantification agreement between meta-analysis (*FC_meta_*) and original constituent studies (*FC_i_*), we computed a MAD-like gene-wise convergence score (Fig. 2a). For each gene *G*:

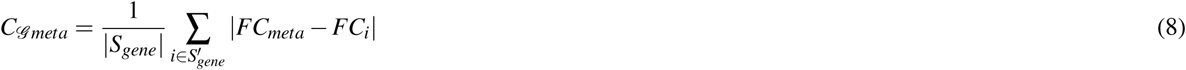

where *S_gene_*and *S’_gene_* denote the valid study set and corresponding indices for the gene (defined in “Overview of the AWmeta framework” section), *S_gene_* the cardinality of the set *S_gene_*, *FC_meta_* = **M** from Eq. 7 and *FC_i_* is the study-exclusive log_2_-based fold change. A lower *C_G meta_* implies better agreement between the meta-analysis and original study estimates within *S_gene_*. For baseline comparison, we also computed, for each original Study *_j_* ∈ *S_gene_*, a gene-wise convergence score:

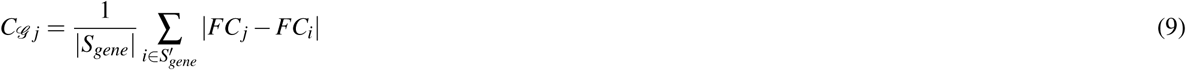

which represents the MAD of Study *_j_*’s fold change from all other contributing studies. This internal consistency benchmark enables direct contrast of AWmeta’s and REM’s convergence performance against the inherent agreement among the original datasets. All comparisons used one-tailed Mann–Whitney test against AWmeta.

### Study-wise convergence assessment for gene differential quantification

To rigorously evaluate the consistency between gene lists derived from meta-analysis methods and those from the original studies, we employed three complementary approaches. For all metrics, higher scores indicate superior study-level convergence.

1. **Adjusted rankeD genE (DE) list similarity**: Our first approach quantifies concordance using a rank-sensitive similarity metric that is critically weighted towards top-ranked genes (Fig. 2g and Extended Data Fig. 7). To construct robustly ordered gene lists (*G*_meta_ for the meta-analysis; *G_i_* for Study*_i_*), we first devised a composite rank for each gene by multiplying its *P*-value rank (ascending) with its |log_2_FC| rank (descending), thereby integrating statistical significance and effect size.

The weighted similarity *S*(*G_meta_, G_i_*) between the meta-analysis and each original study gene lists (containing *N* genes) was computed using a non-linear weighting scheme^91^, which emphasizes the top-ranked gene concordance:

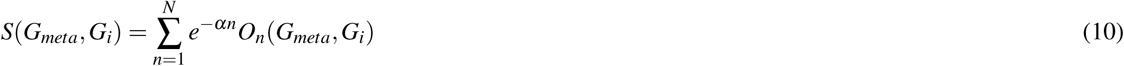

where *O_n_*(*G_meta_, G_i_*) is the number of common genes in the top *n* positions and *α* is a weighting exponent (0.001). This score was then normalized to the interval [−1, 1]^22^ yielding the adjusted similarity:

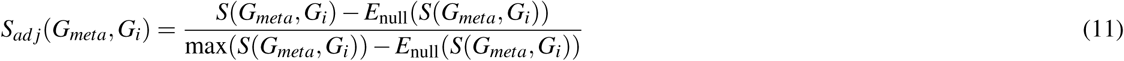

where 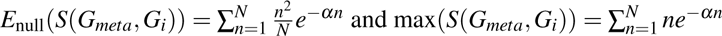 are the expected and maximum scores under a null hypothesis of random gene lists.

1. **Set-based overlap similarity**: To circumvent the limitations of the above rank-dependent approach, which is sensitive to gene ranking variations while potentially overlooking consistent differential expression patterns, we assessed study-wise convergence using a set-based overlap metric that exclusively evaluates binary DEG classification concordance (Fig. 2h,i). Here, DEG sets were determined for both the meta-analysis (*Set_meta_*) and individual studies (*Set_i_*) using predefined statistical thresholds, thereby focusing analytical power on reproducible differential expression status irrespective of positional gene rankings.

We calculated two complementary metrics: 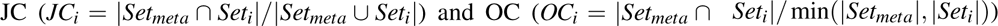. The convergence metric for the meta-analysis relative to Study*_i_* was the arithmetic mean (*JC_i_*+ *OC_i_*)*/*2.

1. **Phi coefficient similarity**: Finally, we measured the association between DEG classifications using phi coefficient^24^ (*φ*) (Fig. 2h,j). This approach considers the extreme case where shared DEGs or non-DEGs between two gene sets might be randomly generated.

For each comparison between the meta-analysis and an original Study*_i_*, we constructed a 2×2 contingency table categorizing all genes as DEG or non-DEG in both datasets and the phi coefficient (*φ_i_*) was then calculated as:

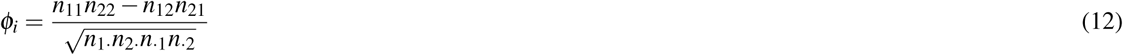

where *n*_11_ represents DEGs, *n*_22_ non-DEGs in both datasets, and *n*_12_ and *n*_21_ represent exclusively-classified DEGs for the binary datasets. The row and column sums are denoted by *n*_1·_, *n*_2·_, *n*_·1_, and *n*_·2_.

To establish a performance baseline, we computed all three convergence metrics for every pairwise combination of the original studies. Overall study-wise convergence differences among AWmeta, REM and baselines were tested by Kruskal–Wallis test, followed by Nemenyi post-hoc test for pairwise comparisons.

For the set-based and *φ* metrics, which rely on binary DEG and non-DEG classification, we note that these outcomes are mutually exclusive and complementary, and therefore report the results derived from the DEG sets for clarity and conciseness.

### Stability and robustness assessment of transcriptomic integration

To demonstrate AWmeta’s resilience, we evaluated both stability—against stochastic sampling—and robustness—against dataset perturbations—using the adjusted DE list similarity (“Study-wise convergence assessment for gene differential quantification” section and Extended Data Fig. 7).

1. **Within-study subsampling stability**: For each original cohort, we randomly partitioned case and control samples of every study into two equal subcohorts, yielding paired “half-study” datasets. Each half-study set underwent independent DEG analysis and subsequent meta-analysis. The similarity between the resulting ordered gene lists was computed over 100 bootstrap replicates, quantifying stability under within-study sampling (Fig. 4a). AWmeta and REM stability distributions were compared via one-tailed Welch’s *t*-test.
2. **External robustness**: We assessed resilience to new data by sequentially incorporating one independent external study (from a held-out pool) into the original meta-analysis (Fig. 4c). For each addition, we performed meta-analysis pre- and post-inclusion, then computed adjusted DE list similarity between resulting ordered gene lists, measuring the impact of disparate external data (AWmeta versus REM, one-tailed Mann–Whitney test).
3. **Internal robustness**: We evaluated sensitivity to study omission by performing leave-one-study-out analyses (Fig. 4e): each original study was removed in turn, and meta-analyses were rerun on the reduced datasets. The similarity between each leave-one-study-out and the full-cohort ranked gene lists, across all iterations, quantified internal robustness (AWmeta versus REM, one-tailed Mann–Whitney test).

### Biological relevance assessment of gene differential quantification

To quantify disease-context relevance of gene differential quantification, we assembled benchmark gene sets for Parkinson’s and Crohn’s disease from three sources: (1) DisGeNET with gene-disease association (GDA) score *>* 0.2^92,93^, (2) MalaCards, and (3) our in-house curated disease-related genetic variation corpus. For reference comparisons, original study-derived biological relevance results serve as baselines.

For each method (AWmeta, REM or baselines), all analyzed genes were ranked twice—(i) by descending |log_2_FC|, (ii) by ascending *P*-value—then each benchmark gene’s ranks were multiplied:

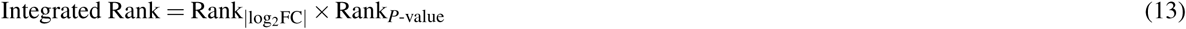

Benchmark genes were then re-ranked according to this Integrated Rank (ascending) to obtain Rank_Integrated Rank_. The Biological Relevance score was calculated for each benchmark gene as:

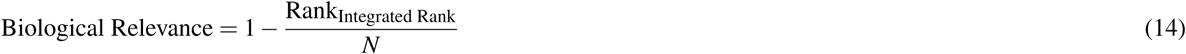

where *N* is the size of gene list from AWmeta, REM or baselines. Higher scores reflect greater biological relevance, signifying that benchmark genes attain superior rankings through combining statistical significance and fold change (Fig. 5a). This rank-based score accounts for gene list size heterogeneity and avoids arbitrary DEG thresholds. We compared biological relevance distributions from AWmeta, REM and baselines using Kruskal-Wallis and Nemenyi post-hoc test.

## Data availability

All transcriptomic datasets used in this study are publicly available via GEO, SRA, and ArrayExpress. Parkinson’s disease data include substantia nigra (GEO accessions: `GSE114517`, `GSE8397`, `GSE20163`, `GSE20164`, `GSE20292`, `GSE7621`, `GSE49036`, `GSE42966`, `GSE43490`, `GSE26927`, `GSE54282`) and peripheral blood (GEO accessions: `GSE57475`, `GSE54536`, `GSE34287`, `GSE99039`, `GSE72267`, `GSE6613`, `GSE18838`, `GSE165082`). Crohn’s disease datasets include peripheral blood (GEO accessions: `GSE119600`, `GSE112057`, `GSE94648`), ileal mucosa (GEO accessions: `GSE102133`, `GSE75214`, `GSE16879`, `GSE68570`, `GSE101794`, `GSE57945`), and colonic mucosa (GEO accessions: `GSE75214`, `GSE16879`, `GSE36807`, `GSE4183`, `GSE9686`, `GSE66207`; ArrayExpress accession: `E-MTAB-184`). Benchmark gene sets for both diseases were retrieved from DisGeNET (https://www.disgenet.com/) and MalaCards (https://www.malacards.org/).

## Code availability

AWmeta is available on GitHub at https://github.com/YanshiHu/AWmeta.

## Acknowledgements

The authors thank all members from Ming Chen’s Group of Bioinformatics for insightful discussions. This work was supported by the National Key Research and Development Program of China (2023YFE0112300); National Natural Sciences Foundation of China (32261133526; 32270709; 32070677); the 151 talent project of Zhejiang Province (first level), the Science and Technology Innovation Leading Scientist (2022R52035).

## Author contributions

Y.S.H. conceptualized the study. M.C. supervised the study. Y.S.H., Z.W., and Y.H. performed data analysis. Y.S.H., Z.W., C.F., and Q.F. contributed to writing the manuscript. All authors reviewed and approved the final manuscript.

## Publisher’s note

Springer Nature remains neutral with regard to jurisdictional claims in published maps and institutional affiliations.

---

**Extended Data Fig. 1.**
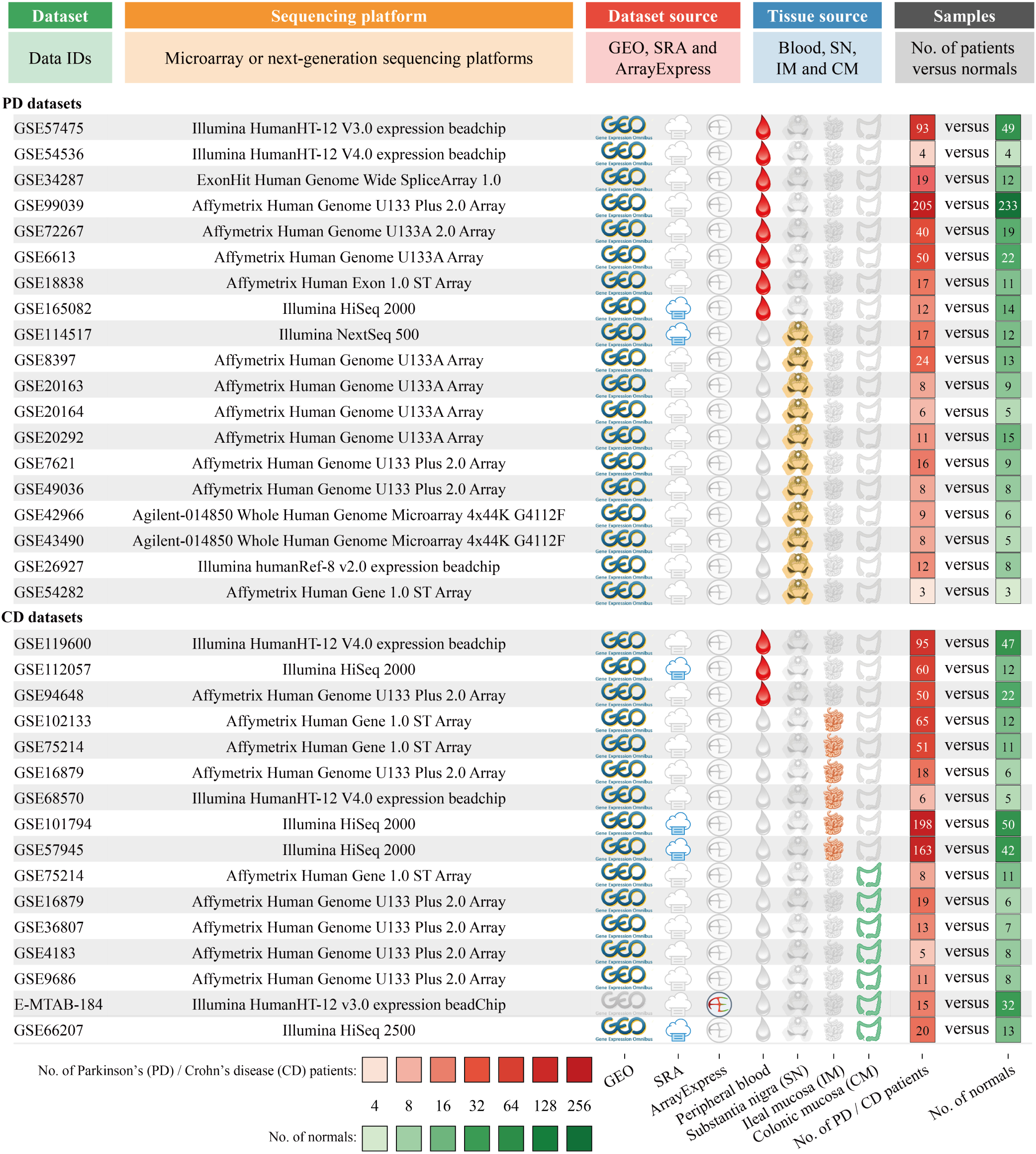
Overview of transcriptomic datasets for Parkinson’s and Crohn’s disease. Datasets for Parkinson’s disease (PD) include substantia nigra^53–60^ and peripheral blood^56,61–68^. Crohn’s disease (CD) datasets comprise ileal mu-cosa^72–77^, colonic mucosa^76–82^, and peripheral blood^69–71^. Detailed metadata include data accession IDs, sequencing platform identifiers, dataset and tissue sources, and patient and control sample sizes. The following icons represent different tissue sources: 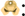 substantia nigra (SN); 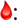 peripheral blood; 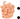 ileal mucosa (IM); 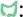 colonic mucosa (CM).

**Extended Data Fig. 2.**
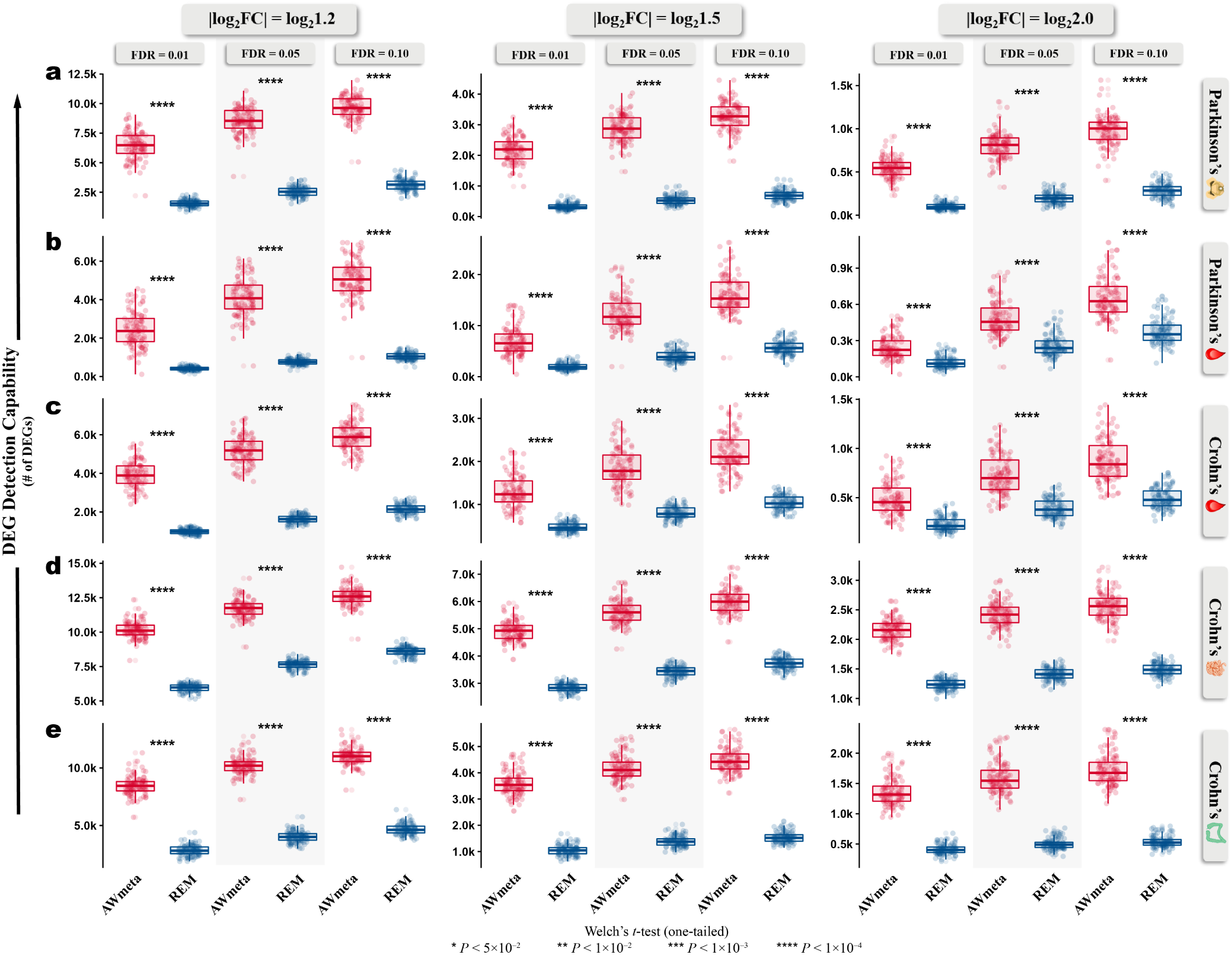
DEG detection capability evaluation across five disease tissues and diverse thresholds. **a**–**e**, DEG detection capability (the number of identified DEGs) was assessed against AWmeta and REM using nine distinct thresholds, combining three corrected *P*-values (FDR) (0.01, 0.05, and 0.10) and three log_2_-based fold change (|log_2_FC|) cutoffs (log_2_1.2, log_2_1.5, and log_2_2.0), spanning Parkinson’s substantia nigra (**a**), Parkinson’s (**b**) and Crohn’s (**c**) peripheral blood, and Crohn’s ileal (**d**) and colonic (**e**) mucosa. Detailed description of DEG detection capability can be referred to in “DEG detection capability evaluation” section in Methods and Fig. 1b. Statistical significance was determined using one-tailed Welch’s *t*-test. Boxplot bounds show interquartile ranges (IQR), centers indicate median values, and whiskers extend to 1.5 IQR. The following icons represent different tissue sources: 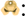 substantia nigra; 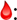 peripheral blood; 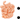 ileal mucosa; 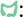 colonic mucosa.

**Extended Data Fig. 3.**
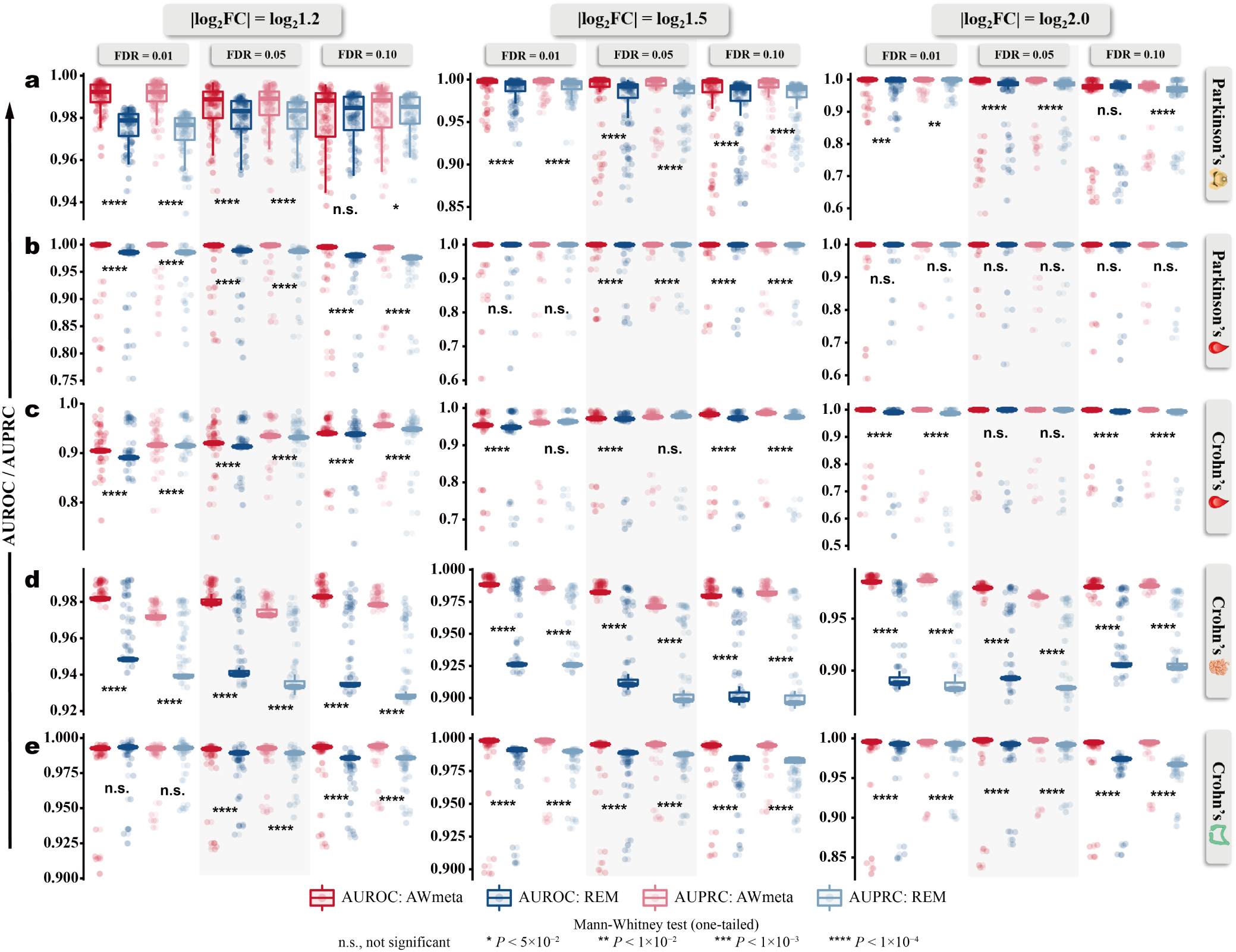
DEG discrimination evaluation across five disease tissues using minimum-permuted semi-synthetic simulation strategy. **a**–**e**, DEG discrimination (AUROC and AUPRC) was assessed against AWmeta and REM using nine distinct thresholds, combining three corrected *P*-values (FDR) (0.01, 0.05, and 0.10) and three log_2_-based fold change (|log_2_FC|) cutoffs (log_2_1.2, log_2_1.5, and log_2_2.0), spanning Parkinson’s substantia nigra (**a**), Parkinson’s (**b**) and Crohn’s (**c**) peripheral blood, and Crohn’s ileal (**d**) and colonic (**e**) mucosa. Detailed description of minimum-permuted semi-synthetic simulation strategy can be referred to in “DEG discrimination evaluation using semi-synthetic simulation strategy” section in Methods, “AWmeta secures robust higher-fidelity DEG identification across transcriptomic contexts” section in Results and Fig. 1d–f. Statistical significance was determined using one-tailed Mann-Whitney test. Boxplot bounds show interquartile ranges (IQR), centers indicate median values, and whiskers extend to 1.5×IQR. The following icons represent different tissue sources: 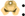 substantia nigra; 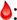 peripheral blood; 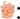 ileal mucosa; 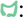 colonic mucosa.

**Extended Data Fig. 4.**
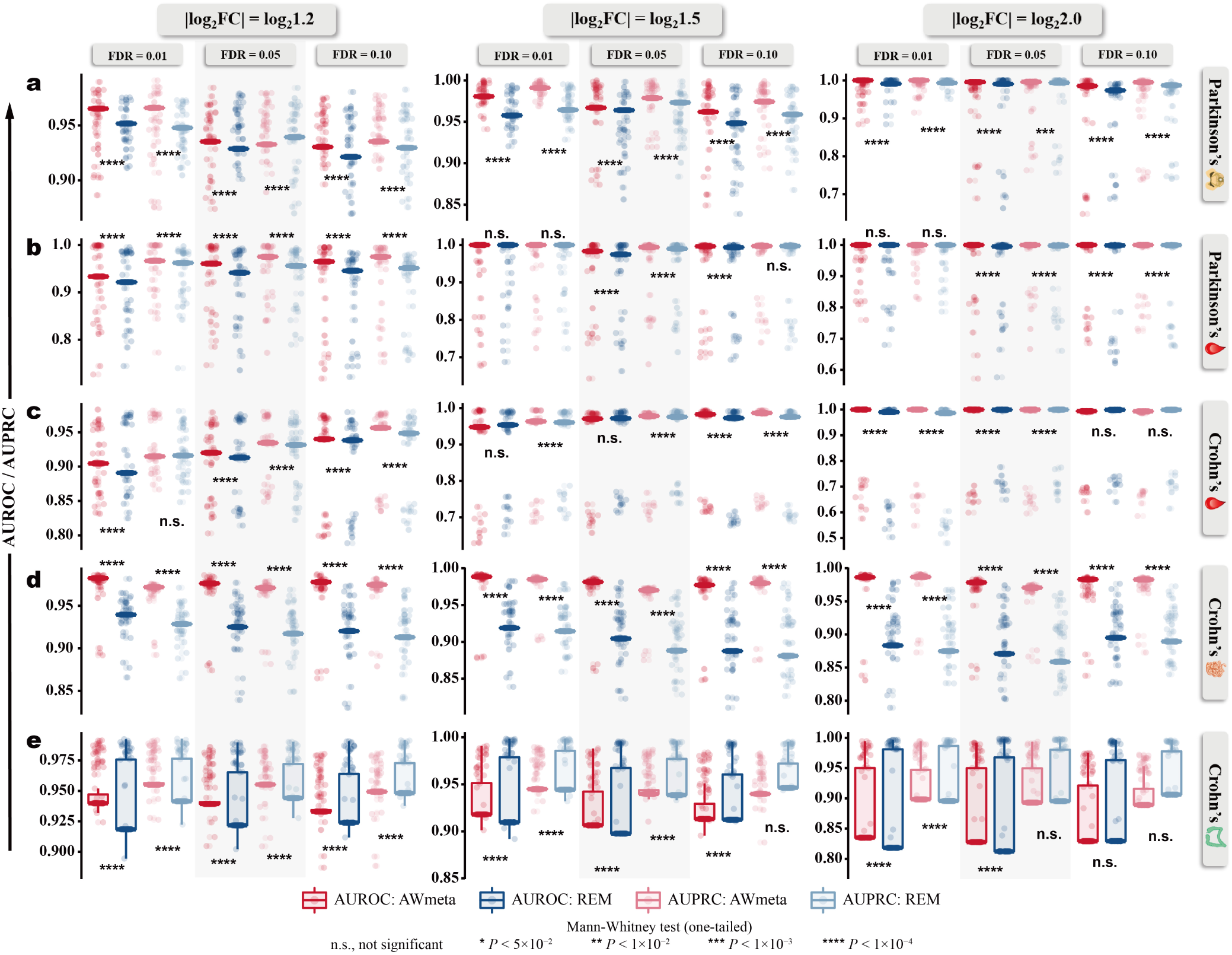
DEG discrimination evaluation across five disease tissues using median-permuted semi-synthetic simulation strategy. **a**–**e**, DEG discrimination (AUROC and AUPRC) was assessed against AWmeta and REM using nine distinct thresholds, combining three corrected *P*-values (FDR) (0.01, 0.05, and 0.10) and three log_2_-based fold change (|log_2_FC|) cutoffs (log_2_1.2, log_2_1.5, and log_2_2.0), spanning Parkinson’s substantia nigra (**a**), Parkinson’s (**b**) and Crohn’s (**c**) peripheral blood, and Crohn’s ileal (**d**) and colonic (**e**) mucosa. Detailed description of median-permuted semi-synthetic simulation strategy can be referred to in “DEG discrimination evaluation using semi-synthetic simulation strategy” section in Methods, “AWmeta secures robust higher-fidelity DEG identification across transcriptomic contexts” section in Results and Fig. 1d–f. Statistical significance was determined using one-tailed Mann-Whitney test. Boxplot bounds show interquartile ranges (IQR), centers indicate median values, and whiskers extend to 1.5×IQR. The following icons represent different tissue sources: 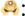 substantia nigra; 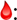 peripheral blood; 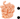 ileal mucosa; 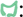 colonic mucosa.

**Extended Data Fig. 5.**
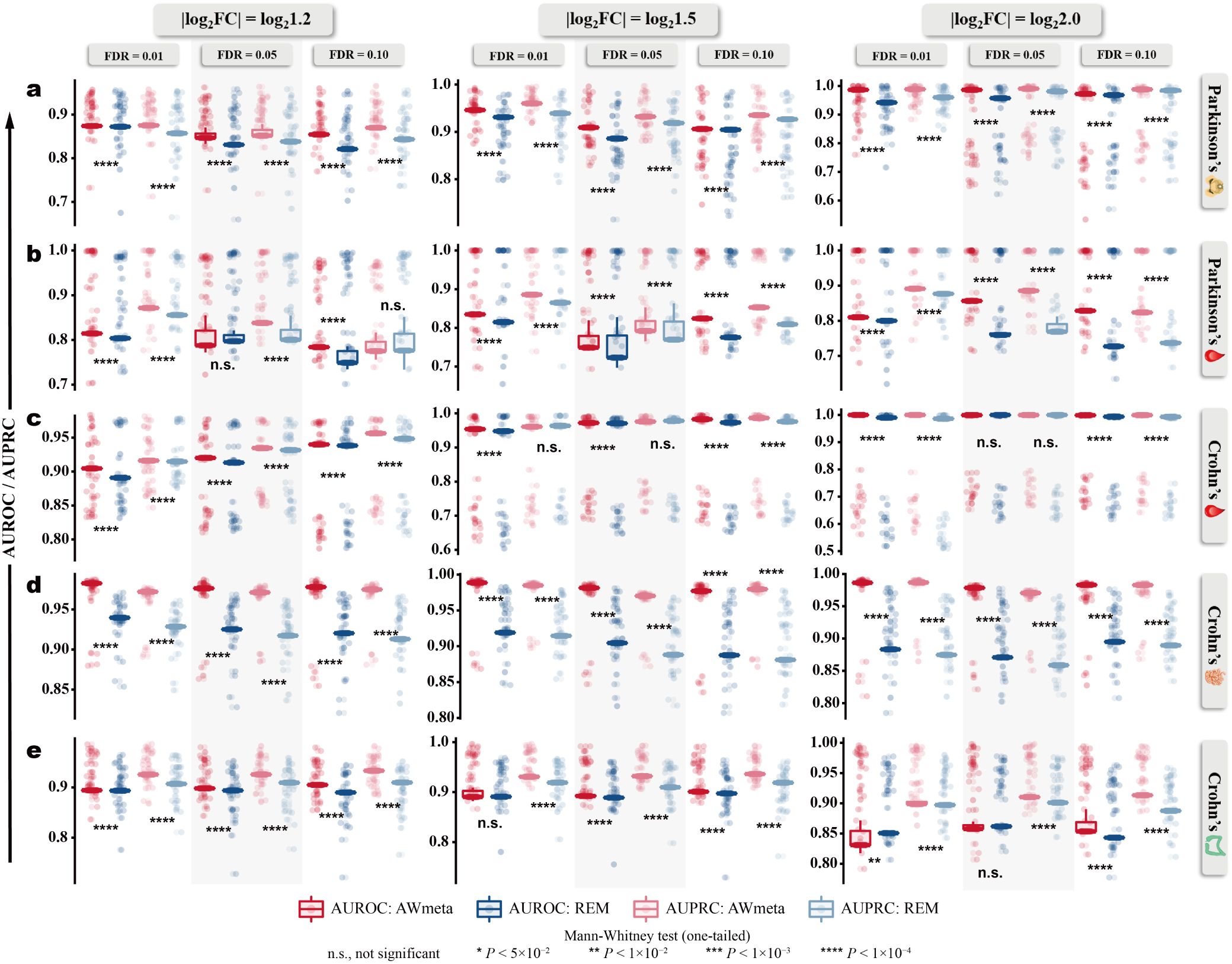
DEG discrimination evaluation across five disease tissues using maximum-permuted semi-synthetic simulation strategy. **a**–**e**, DEG discrimination (AUROC and AUPRC) was assessed against AWmeta and REM using nine distinct thresholds, combining three corrected *P*-values (FDR) (0.01, 0.05, and 0.10) and three log_2_-based fold change (|log_2_FC|) cutoffs (log_2_1.2, log_2_1.5, and log_2_2.0), spanning Parkinson’s substantia nigra (**a**), Parkinson’s (**b**) and Crohn’s (**c**) peripheral blood, and Crohn’s ileal (**d**) and colonic (**e**) mucosa. Detailed description of maximum-permuted semi-synthetic simulation strategy can be referred to in “DEG discrimination evaluation using semi-synthetic simulation strategy” section in Methods, “AWmeta secures robust higher-fidelity DEG identification across transcriptomic contexts” section in Results and Fig. 1d–f. Statistical significance was determined using one-tailed Mann-Whitney test. Boxplot bounds show interquartile ranges (IQR), centers indicate median values, and whiskers extend to 1.5×IQR. The following icons represent different tissue sources: 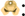 substantia nigra; 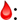 peripheral blood; 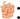 ileal mucosa; 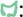 colonic mucosa.

**Extended Data Fig. 6.**
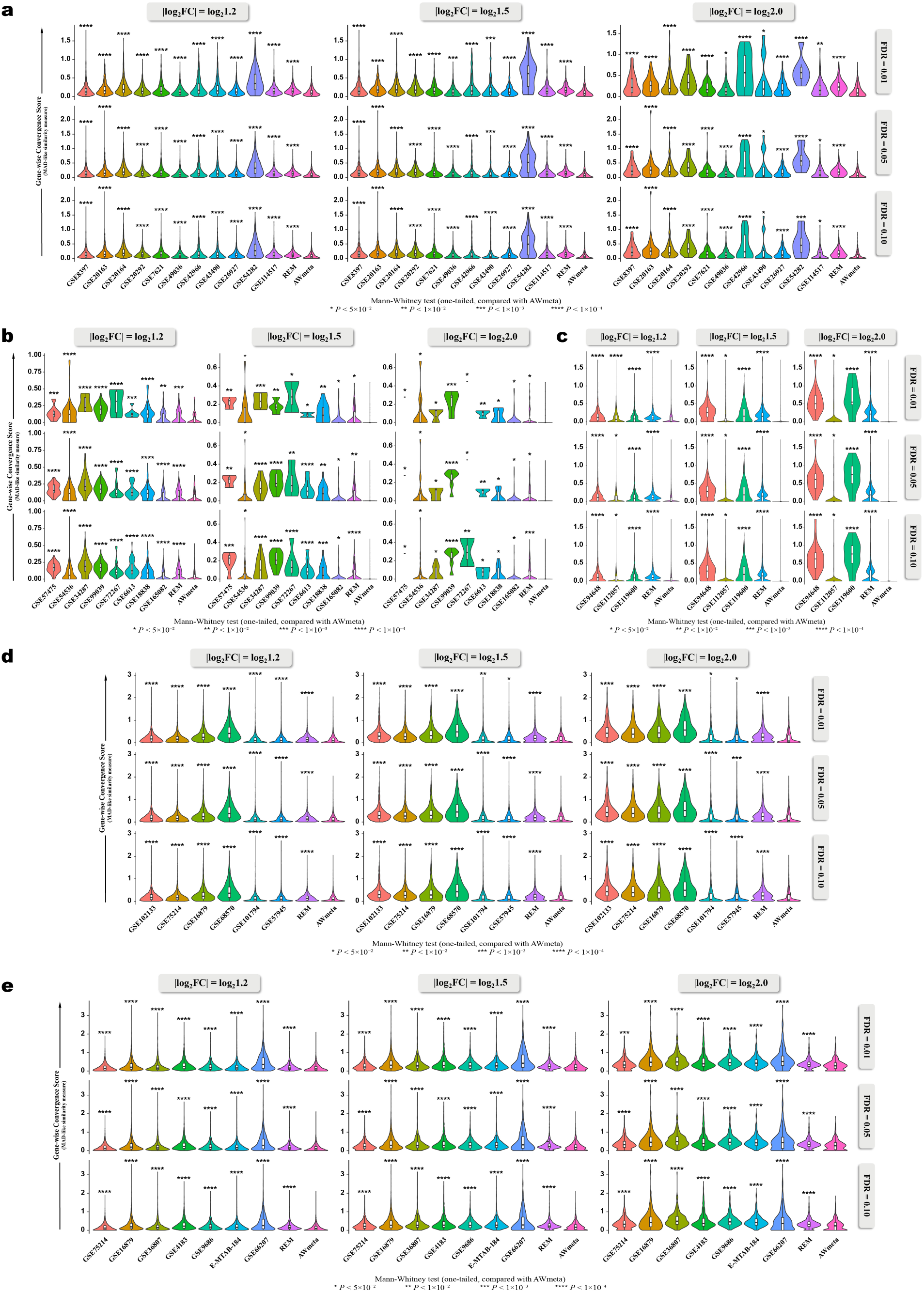
AWmeta establishes superior DEG-wise convergence in gene differential quantification. **a**–**e**, Considering that DEGs instead of non-DEGs are primarily involved in disease etiology, to explore whether gene-(unfiltered) and DEG-wise convergence assessment results are different, mean absolute deviation (MAD)-like similarity measure was utilized to quantify the per-DEG fold change (|log_2_FC|) similarity among AWmeta, REM and original studies, with smaller values indicating better convergence, which demonstrates AWmeta’s consistent superior DEG-with gene-wise convergence over REM and orignal studies using nine distinct thresholds, combining three corrected *P*-values (FDR) (0.01, 0.05, and 0.10) and three log_2_-based fold change (|log_2_FC|) cutoffs (log_2_1.2, log_2_1.5, and log_2_2.0), across five disease tissues: Parkinson’s substantia nigra (**a**), Parkinson’s (**b**) and Crohn’s (**c**) peripheral blood, and Crohn’s ileal (**d**) and colonic (**e**) mucosa. For comparison purpose, results from original studies serve as reference baselines. Statistical significance of REM and baselines against AWmeta for DEG-wise convergence comparisons was tested with one-tailed Mann-Whitney test. Detailed description for MAD-like DEG-wise convergence similarity measure can be referred to in “Gene-wise convergence assessment for gene differential quantification” section in Methods, “AWmeta establishes superior gene- and study-wise convergence in gene differential quantification” section in Results and Fig. 2a. Boxplot bounds show interquartile ranges (IQR), centers indicate median values, and whiskers extend to 1.5×IQR. The following icons represent different tissue sources: 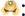 substantia nigra; 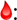 peripheral blood; 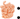 ileal mucosa; 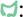 colonic mucosa.

**Extended Data Fig. 7.**
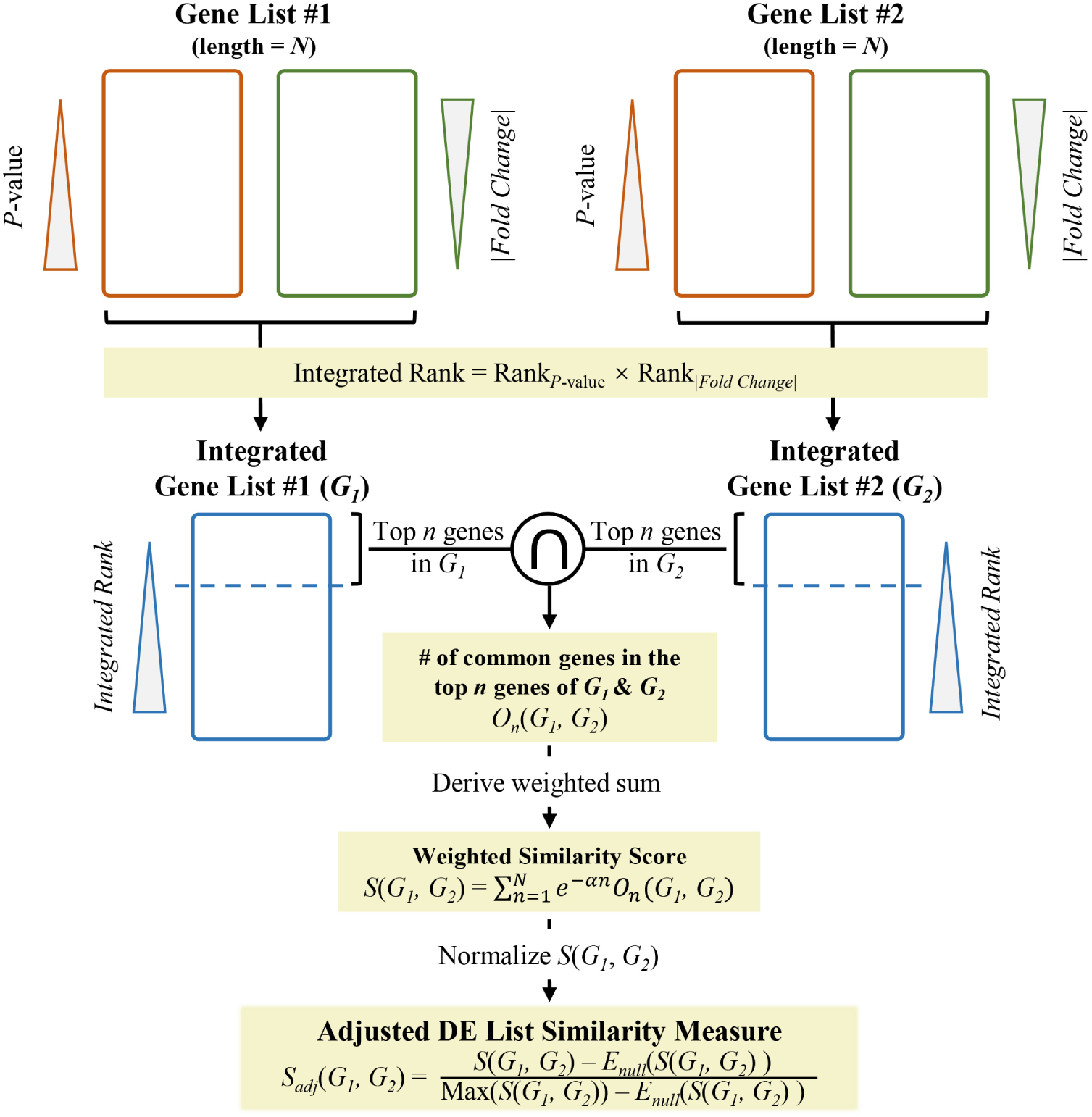
Conceptual schematic of adjusted DE list similarity measure. The methodological details reside in “Study-wise convergence assessment for gene differential quantification” section in Methods.

**Extended Data Fig. 8.**
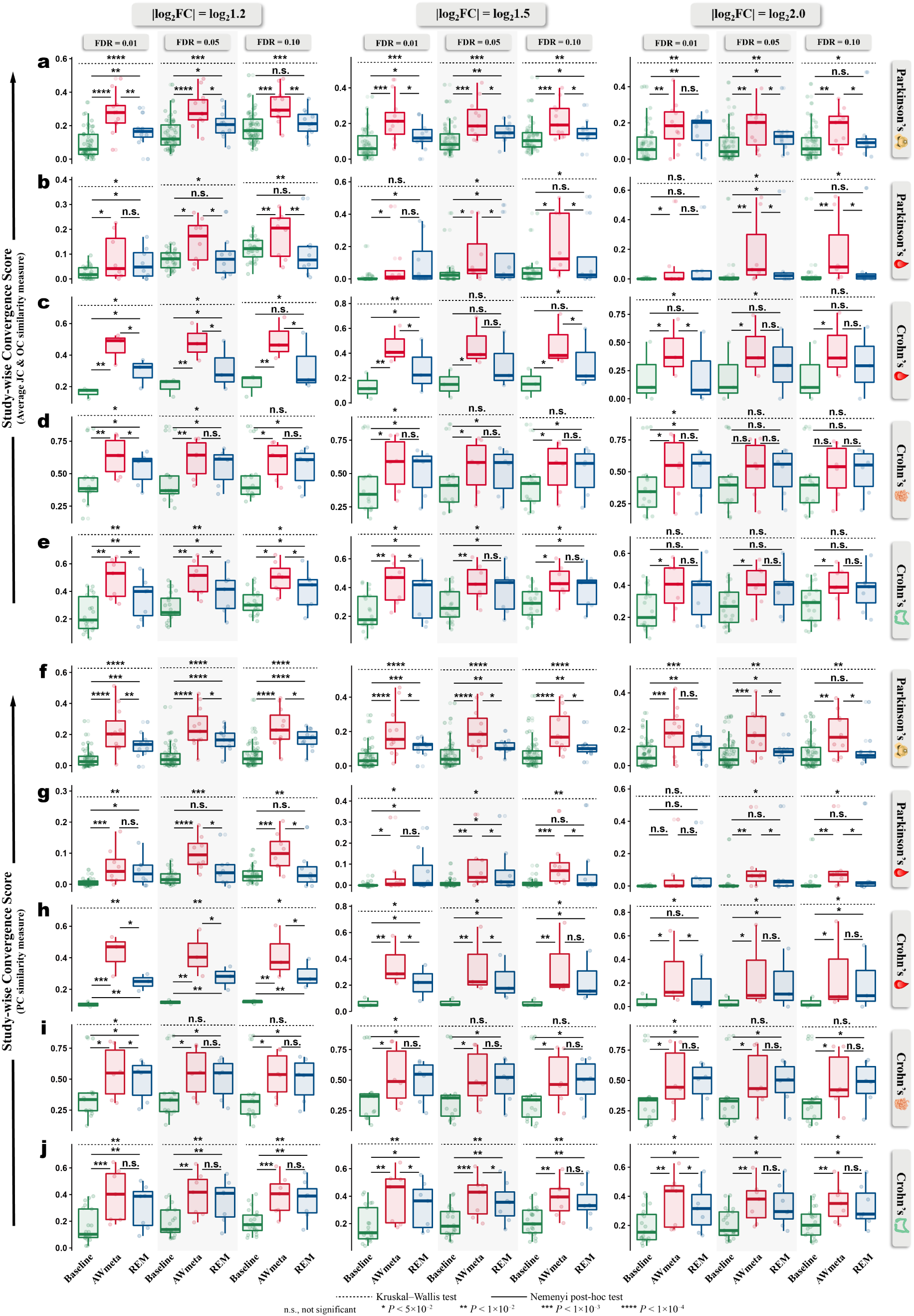
AWmeta maintains robust superior study-wise convergence in gene differential quantification across diverse thresholds. **a**–**j**, To dissect whether set-theory-based similarity-derived study-wise convergence assessment results vary with diverse DEG thresholds, we used nine distinct thresholds, combining three corrected *P*-values (FDR) (0.01, 0.05, and 0.10) and three log_2_-based fold change (|log_2_FC|) cutoffs (log_2_1.2, log_2_1.5, and log_2_2.0), to benchmark study-wise convergence, which showcases AWmeta maintains robust superior study-wise convergence in gene differential quantification across diverse thresholds over REM and baselines in five disease tissues, both by means of the average of Jaccard (JC) and overlap coefficient (OC) (**a**–**e**) and phi coefficient (PC) (**f**–**j**). For comparison purpose, results from original studies serve as reference baselines. Overall study-wise convergence differences among AWmeta, REM and baselines were tested by Kruskal–Wallis test, followed by Nemenyi post-hoc test for pairwise comparisons. Details for these two study-wise convergence similarity measures appear in “Study-wise convergence assessment for gene differential quantification” section in Methods and Fig. 2h–j. Boxplot bounds show interquartile ranges (IQR), centers indicate median values, and whiskers extend to 1.5×IQR. The following icons represent different tissue sources: 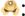 substantia nigra; 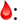 Peripheral blood; 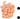 ileal mucosa; 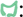 colonic mucosa.

